# Hand2 delineates mesothelium progenitors and is reactivated in mesothelioma

**DOI:** 10.1101/2020.11.11.355693

**Authors:** Karin D. Prummel, Helena L. Crowell, Susan Nieuwenhuize, Eline C. Brombacher, Stephan Daetwyler, Charlotte Soneson, Jelena Kresoja-Rakic, Manuel Ronner, Agnese Kocere, Alexander Ernst, Zahra Labbaf, David E. Clouthier, Anthony B. Firulli, Héctor Sánchez-Iranzo, Sundar R. Naganathan, Rebecca O’Rourke, Erez Raz, Nadia Mercader, Alexa Burger, Emanuela Felley-Bosco, Jan Huisken, Mark D. Robinson, Christian Mosimann

## Abstract

The mesothelium forms epithelial membranes that line the bodies cavities and surround the internal organs. Mesothelia widely contribute to organ homeostasis and regeneration, and their dysregulation can result in congenital anomalies of the viscera, ventral wall defects, and mesothelioma tumors. Nonetheless, the embryonic ontogeny and developmental regulation of mesothelium formation has remained uncharted. Here, we combine genetic lineage tracing, *in toto* live imaging, and single-cell transcriptomics in zebrafish to track mesothelial progenitor origins from the lateral plate mesoderm (LPM). Our single-cell analysis uncovers a post-gastrulation gene expression signature centered on *hand2* that delineates distinct progenitor populations within the forming LPM. Combining gene expression analysis and imaging of transgenic reporter zebrafish embryos, we chart the origin of mesothelial progenitors to the lateral-most, *hand2*-expressing LPM and confirm evolutionary conservation in mouse. Our time-lapse imaging of transgenic *hand2* reporter embryos captures zebrafish mesothelium formation, documenting the coordinated cell movements that form pericardium and visceral and parietal peritoneum. We establish that the primordial germ cells migrate associated with the forming mesothelium as ventral migration boundary. Functionally, *hand2* mutants fail to close the ventral mesothelium due to perturbed migration of mesothelium progenitors. Analyzing mouse and human mesothelioma tumors hypothesized to emerge from transformed mesothelium, we find *de novo* expression of LPM-associated transcription factors, and in particular of Hand2, indicating the re-initiation of a developmental transcriptional program in mesothelioma. Taken together, our work outlines a genetic and developmental signature of mesothelial origins centered around Hand2, contributing to our understanding of mesothelial pathologies and mesothelioma.

## Introduction

As a key feature of the vertebrate body plan, the mesothelium is composed of several continuous, epithelial monolayers surrounding the internal organs (visceral mesothelium) and lining the body cavities (parietal mesothelium). The mesothelium provides a protective layer against invasive microorganisms, produces serous fluid that both decreases friction of moving organs, and enables the transport of cells and nutrients across serosal cavities (Mutsaers, 2002; Mutsaers and Wilkosz, 2007). Moreover, cell tracking studies have established that the mesothelium contributes to a multitude of downstream cell fates including smooth muscles and fibroblasts during organogenesis, tissue homeostasis, and regeneration (Carmona et al., 2018; Koopmans and Rinkevich, 2018; Rinkevich et al., 2012). While mesothelium-lined body cavities are a fundamental trait across bilaterian animals (Hartenstein and Mandal, 2006; Monahan-Earley et al., 2013; Technau and Scholz, 2003), little is known about how the vertebrate mesothelium lineage initially emerges during development.

The embryonic mesothelium, also called coelomic epithelium after establishing its baso-apical polarization, is a highly dynamic cell layer that undergoes epithelial-to-mesenchymal transition (EMT) during development and seeds mesenchymal cells to underlying tissues (Carmona et al., 2018; Rinkevich et al., 2012). Several prior observations tie mesothelial lineage origins to the lateral plate mesoderm (LPM) (Ahn et al., 2002; Carmona et al., 2016; Winters et al., 2012), a mesodermal progenitor territory that forms at the periphery of the early vertebrate embryo (Prummel et al., 2020). In vertebrates, the coelomic cavity forms stereotypically by splitting the LPM into dorsal and ventral layers (Onimaru et al., 2011). A subset of cells within both layers differentiates into polarized epithelial cells that form visceral (splanchnic) mesothelial layers and parietal (somatic) mesothelial layers (Funayama et al., 1999; Sadler and Feldkamp, 2008; Sheng, 2015). Ultimately, the coelom in amniotes spans from the neck to the abdomen and outlines four main body compartments: two pleural cavities (around the lungs), a pericardiac cavity (around the heart), and a peritoneal (abdominal) cavity, each with their associated mesothelial layers (**Fig. 1A**). Representative for teleosts, zebrafish feature mesothelium-lined cardiac and abdominal cavities. Which territories within the emerging LPM initially harbor the mesothelial progenitors, how and when the mesothelium diverges from other LPM lineages, and how the visceral and parietal layers form remain uncharted.

**Figure 1:**
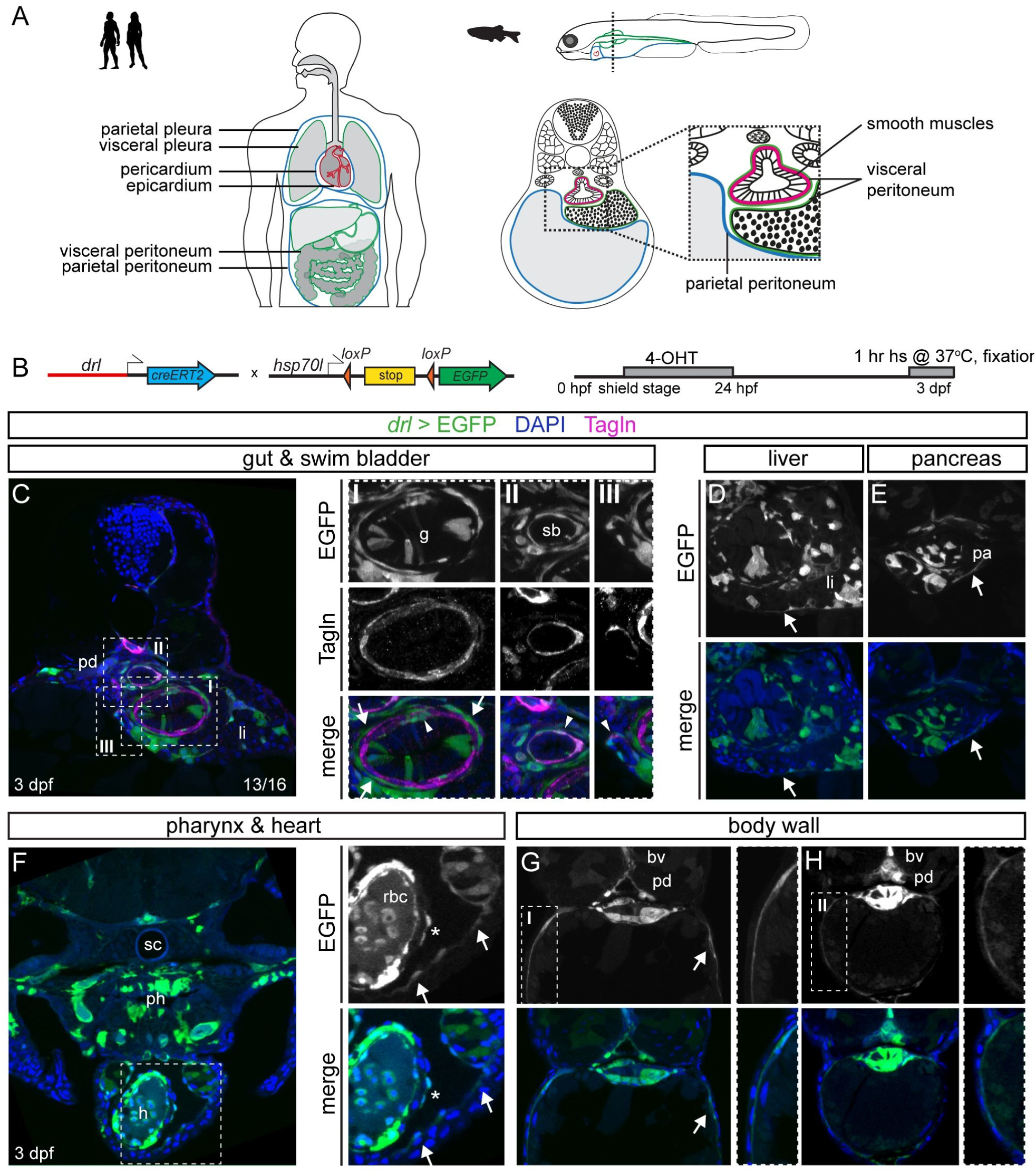
Visceral and parietal mesothelial layers in zebrafish are LPM lineages. **(A)** Mesothelium in human versus zebrafish embryo. Transverse schematic of zebrafish embryo of liver, gut, and budding swim bladder with associated mesothelium and smooth muscle at 3 dpf. **(B)** Tracing LPM using *drl:creERT2* x *hsp70l:Switch*, 4-OHT administered at shield stage and washed off before 24 hpf. *drl>EGFP* indicates LPM lineage labeling. **(C)** Trunk section of *drl* lineage-traced 3 dpf embryo co-stained for smooth muscle (Tagln). Boxed regions show details of EGFP lineage- and smooth muscle-labeling around gut (box I), swim bladder (box II), and liver ducts (box III). Arrows depict Tagln-negative;EGFP-positive cells, arrowheads depict Tagln;EGFP double-positive cells. **(D,E)** EGFP-based LPM labeling in the peritoneum around liver (D) and pancreas (E). **(F)** Rostral transverse section, lineage labeling of pericardium, ventricle (endocardium, myocardium, potentially epicardium (asterisk), blood), and LPM- and endoderm-derived organs in the head (head cartilage, vasculature, pharynx). **(G,H)** Sections of two regions along the anterior-posterior axis, (G) at yolk, (H) at yolk extension, showing *drl*-based LPM lineage labeling of parietal peritoneum forming body wall together with skin layer (boxed region). Pronephric duct (pd), liver (li), gut (g), swim bladder (sb), pancreas (pa), heart (h), blood vessel (bv), spinal cord (sc), pharynx (ph), red blood cells (rbc). Nuclei in blue (DAPI).

Embryonic studies of the mesothelium have predominantly focused on developmental stages after the coelomic epithelium has formed. Several genes, including *Mesothelin* (*Msln*), *Gata4, Tbx18, Tcf21*, and *Wilms Tumor 1* (*Wt1*), have enabled labeling and genetic lineage tracing of mesothelial lineages in mouse and chick (Alghamdi et al., 2020; Ariza et al., 2016; Chau et al., 2014; Delgado et al., 2014; Rinkevich et al., 2012). The *Wt1*-expressing coelomic epithelium contributes to the mature mesothelium, fibroblasts, stellate cells, smooth muscles, and white adipose tissue associated with the gastrointestinal tract, liver, lungs, urogenital system, and heart (Asahina et al., 2011; Cano et al., 2013; Carmona et al., 2016; Delgado et al., 2014; Lee et al., 2019; Lua et al., 2014; Martínez-Estrada et al., 2010; Rinkevich et al., 2012; Sebo et al., 2018; Zhou et al., 2008). Further, regional specification within the mesothelial components along the developing gut is recognizable early in mouse development (Han et al., 2020; Kishimoto et al., 2020). Studies using zebrafish have documented expression of *wt1a/b, tcf21*, and *tbx18* in the epicardium, the visceral mesothelial layer covering the heart (Peralta et al., 2013; Peralta et al., 2014). Despite these advances across models, expression of these conserved genes is initiated after the onset of coelomic epithelium formation, leaving the earliest differentiation steps obscure.

Compromised integrity of the adult mesothelium can result in pathologies including intra-abdominal organ adhesion (Tsai et al., 2018), serosal fibrosis (Mutsaers et al., 2015), pericarditis (Chiabrando et al., 2020), and mesothelioma tumors (Carbone et al., 2019; Yap et al., 2017). Malignant mesothelioma is a rapidly fatal solid tumor that can arise within the visceral or parietal mesothelia, predominantly as the result of environmental exposure to asbestos (Carbone et al., 2019; Felley-Bosco and Macfarlane, 2018; Odgerel et al., 2017; Wagner et al., 1962; Yap et al., 2017). While mesothelioma cases are increasing globally despite regulatory means to curb the use of causative agents, treatment remains limited (Hinz and Heasley, 2019; Yap et al., 2017). Presenting predominantly as epithelioid, sarcomatoid, and biphasic phenotypes, malignant mesothelioma frequently harbor genetic alterations affecting the tumor suppressors *BAP1, NF2, CDKN2AB*, and *TP53* (Bueno et al., 2016; Cheng et al., 1994; Hmeljak et al., 2018; Quetel et al., 2020; Tate et al., 2019). Nonetheless, the cell of origin and the underlying aberrant molecular mechanisms leading to mesothelioma remain uncertain.

In addition to the mesothelium, the LPM forms a vast array of downstream cell fates that include the cardiovascular system, blood, kidneys, and limb connective tissue (Prummel et al., 2020). How the LPM partitions into its diverse fates and what regulatory programs specify the individual progenitor fields remains unclear. Emerging as a dedicated mesendoderm domain, the post-gastrulation LPM segments into recognizable bilateral territories discernible by the expression of several transcription factor genes including *Scl/Tal1, Lmo2, Pax2a, Nkx2.5*, and *Hand1/2* (Chal and Pourquié, 2017; Davidson and Zon, 2004; Prummel et al., 2020; Takasato and Little, 2015). *dHand/Hand2*, encoding a conserved basic helix-loop-helix transcription factor, is expressed during segmentation stages in the most laterally positioned LPM progenitors in amphioxus and zebrafish (Onimaru et al., 2011; Perens et al., 2016; Prummel et al., 2019). In the developing mouse embryo, *Hand2* expression has been described in the flank at comparable embryonic stages (Charité et al., 2000; Firulli et al., 1998; Srivastava et al., 1995). Studies across vertebrate models have revealed key insights into the contribution of Hand2 and its paralog eHand/Hand1 in anterior LPM (ALPM) progenitors that contribute to the heart, (fore)limbs, and branchial arches (Funato et al., 2009; Yelon et al., 2000; Yin et al., 2010). Additionally, in the posterior LPM of zebrafish and chick, Hand2 has been linked to refining the fate divergence between smooth muscle versus hemangioblast and kidney fates during somitogenesis (Gays et al., 2017; Perens et al., 2016; Shin et al., 2009). The definitive fate of especially the posterior Hand2-expressing progenitors has remained unclear.

Here, we establish several lines of evidence that *hand2* in zebrafish, both by expression and by function, is the earliest specific transcription factor gene demarcating the emerging mesothelial progenitors within the LPM. We provide further evidence for conservation of this property in mouse. We link the developmental function of Hand2 in mesothelium formation to a reactivation of an early coelomic epithelium-focused LPM program in mouse and human mesothelioma tumors. Our findings propose that Hand2 expression contributes to the unique properties of mesothelial progenitor cells in development and in mesothelioma.

## Results

### Zebrafish mesothelium is LPM-derived

To formally assess whether in zebrafish mesothelial membranes are *bona fide* LPM lineages, we performed genetic lineage tracing using *drl:creERT2* that is active in LPM-primed progenitor cells from the onset of gastrulation until early-to-mid somitogenesis (Mosimann et al., 2015; Prummel et al., 2019). We induced *drl:creERT2;hsp70l:Switch* embryos with 4-OH-Tamoxifen (4-OHT) at shield stage and analyzed the resulting EGFP-based LPM lineage labeling in transverse sections at 3 days post-fertilization (dpf) when LPM-derived organs are clearly detectable (**Fig. 1A,B**). EGFP expression recapitulated broad labeling of LPM-derived organs including endothelium, blood, and cardiac lineages, and sparse labeling of endoderm-derived organs, in line with previous observations using *drl:creERT2* (Mosimann et al., 2015; Prummel et al., 2019) (**Fig. 1C-G, Fig. S1A-C**). Extending the previously reported LPM lineage labeling in the Transgelin (Tagln)-positive smooth muscle layers around the zebrafish gut (**Fig. 1C**)(Gays et al., 2017), we also observed smooth muscles around the swim bladder and ducts within the liver by double-positive staining for EGFP and Tagln (**Fig. 1C**). In addition, we consistently observed EGFP-positive, yet Tagln-negative cells as thin epithelial layers surrounding the gut and swim bladder (**Fig. 1C**) as well as around other endodermal organs including the liver and the pancreas (**Fig. 1D,E**). We also found EGFP lineage labeling in the pericardial and epicardial layers surrounding the heart, confirming their LPM origin (**Fig. 1F, Fig. S1A,D-I**). Of note, at this stage, the pericardium directly adheres to the body wall, forming a sac around the cardiac cavity, while in adult zebrafish the pericardium establishes a dedicated body cavity. Lastly, we observed prominent *drl*-based LPM lineage labeling within the body wall as an epithelial layer right underneath the skin surrounding the yolk and the yolk extension along the anterior-posterior axis (**Fig. 1G,H**). From these observations, we conclude that the prospective coelomic epithelium and most, if not all, developing mesothelial layers in zebrafish are LPM-derived.

### The early LPM consists of distinct progenitor populations

We next sought to chart the mesothelial progenitors within the emerging LPM. To probe whether early mesothelial progenitors can be recognized within the emerging LPM, we analyzed the transcriptome of individual zebrafish LPM cells by single-cell RNA-sequencing (scRNA-seq) at tailbud stage (**Fig. 2A**). At this stage, the end of gastrulation, the *drl:mCherry* reporter-labeled LPM comprises approximately 7% of all cells in the zebrafish embryo (**Fig. 2B, Fig. S2**). Upon quantifying the transcriptomes from individually sorted *drl:mCherry-*positive cells using CEL-Seq2 (Muraro et al., 2016), we obtained 1039 cells that passed filtering and quality control. Using graph-based clustering (Louvain algorithm (Blondel et al., 2008)), we called 15 distinct cell populations within the *drl*-positive LPM at tailbud stage (**Fig. 2C**). We annotated these as contributing to 8 major subpopulations based on canonical markers and published gene expression patterns in conjunction with marker genes identified through differential expression analysis (**Fig. 2E**, see Methods for details).

**Figure 2:**
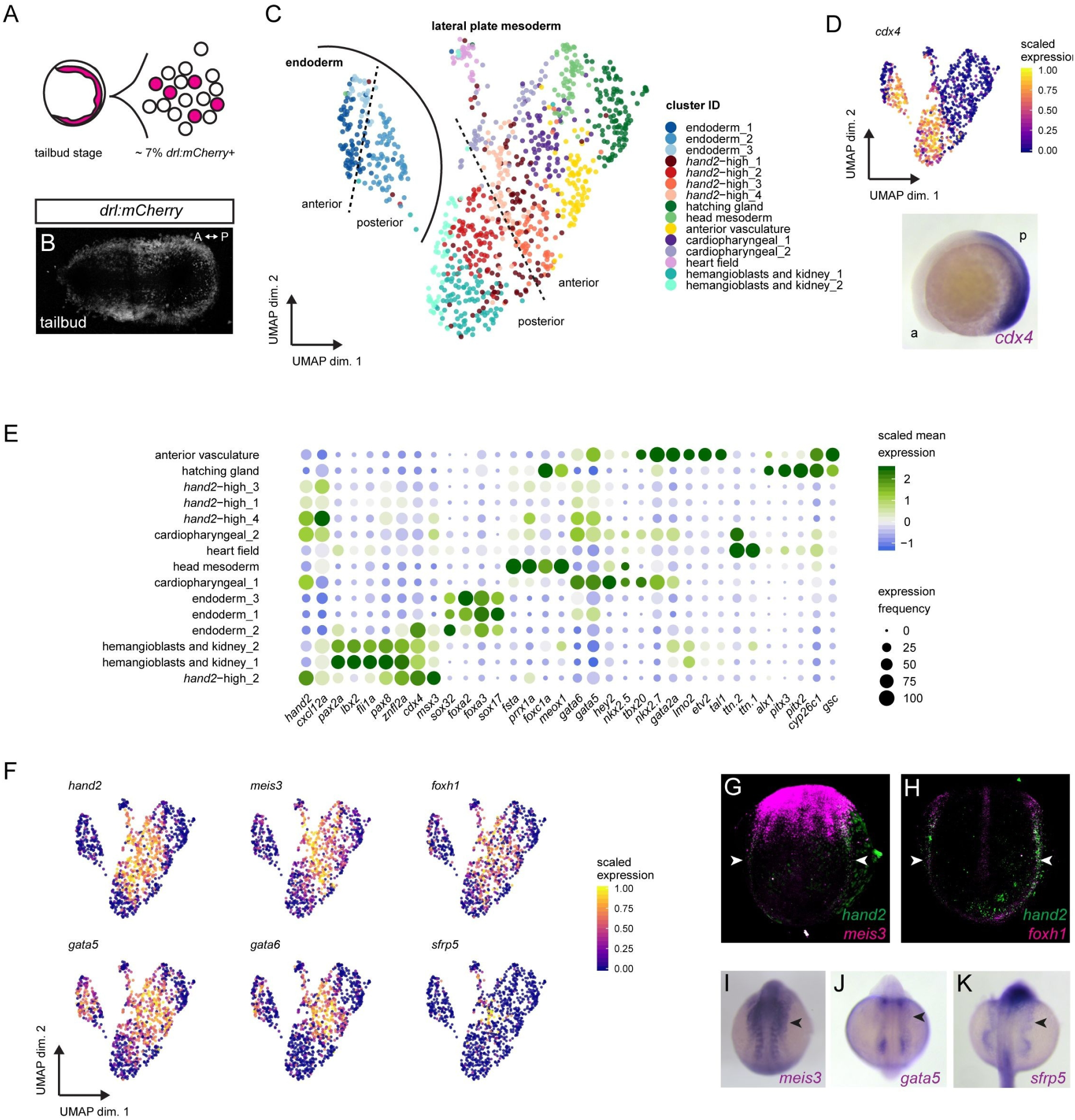
scRNA-sequencing of early LPM reveals a heterogeneous progenitor pool. **(A)** LPM-marking *drl:mCherry*-positive cells of zebrafish embryos at tailbud stage, FACS-isolated, and sequenced using CelSeq2. **(B)** SPIM projection of *drl:mCherry* embryo at tailbud, labeling the LPM. **(C)** UMAP plot revealing 15 LPM cell clusters, colored by subpopulation: cardiopharyngeal (1,2), heart field, anterior vasculature, hemangioblasts/kidney (1,2), head mesoderm, hatching gland, *hand2*-high (1-4), endoderm (1-3) as distinct group. (**D**) Approximate anterior-to-posterior orientation based on *cdx4* (RNA ISH at 5 ss). **(E)** Dotplot including key cell fate markers to annotate clusters. Dots colored by column-scaled mean expression (log-transformed library-size-normalized counts) and sized by expression frequency (fraction of cells with non-zero counts); rows and clusters ordered by hierarchical clustering of scaled expression values. **(F)** UMAP plots of several genes co-expressed with *hand2* or among cluster-determining genes in four *hand2*-high clusters. Cells colored by scaled expression values using lower/upper 1%-quantile boundaries. (**G-K**) Whole-mount gene expression analysis of select transcripts enriched in *hand2*-high cells by fluorescent *in situ* hybridization (**G,H**) and mRNA ISH (**I-K**). Fluorescent ISH of *meis3* (**G**), *foxh1* (**H**) with *hand2* at 4 ss, revealing overlap in posterior LPM. *meis3* ISH at 10 ss (**I**), *gata5* at 12 ss (**J**), and *sfrp5* at 18 ss (**K**) showing expression in lateral-most LPM sprawling outwards (arrowheads).

While mRNA *in situ* hybridization (ISH) and transgene expression for the earliest markers of individual LPM fate potentials renders them chiefly detectable from early somitogenesis on (Davidson and Zon, 2004), our analysis at tailbud stage resolved seemingly determined LPM progenitor fields already at the end of gastrulation, in line with and extending previous findings in zebrafish and mouse embryos (Farrell et al., 2018; Mohammed et al., 2017; Mosimann et al., 2015; Pijuan-Sala et al., 2019; Scialdone et al., 2016; Wagner et al., 2018). Our uncovered clusters broadly represent the cardiopharyngeal, emerging head mesoderm, hatching gland progenitors, endoderm, and endothelial and hematopoietic progenitors (**Fig. 2C,E**). Three clusters were composed of presumptive cardiopharyngeal progenitors based on the expression of *tbx20, nkx2.5, nkx2.7, hey2, gata4/5/6, ttn.1*, and *ttn.2* (**Fig. 2C,E**) (Bloomekatz et al., 2017; Gibb et al., 2018; Lee et al., 1996; Lu et al., 2017; Peterkin et al., 2009; Shih et al., 2015; Wang et al., 2019a). One cluster was positive for designated markers of putative head mesoderm progenitors, including *fsta, foxc1a, gsc, meox1*, and *prrx1a* (Wang et al., 2019b). Based on the expression pattern of *alx1, pitx2*, and *pitx3*, we assigned a cluster to represent the hatching gland progenitors (John et al., 2013; Zilinski et al., 2005), in line with *drl* reporter activity in these cells (Mosimann et al., 2015). We additionally uncovered cells expressing the endodermal genes *sox32, sox17, gata5, gata6, foxa2*, and *fox3a* (Farrell et al., 2018; Schier and Talbot, 2001; Wagner et al., 2018; Warga and Nüsslein-Volhard, 1999) as distinct group of clusters within the analyzed cells (**Fig. 2C,E**). This observation is in line with previous findings that the *drl* reporter-positive cells at tailbud stage represent either mixed endoderm- and LPM-primed populations or a bi-potential LPM-fated mesendoderm population (Prummel et al., 2019).

Towards reconstructing rudimentary positional information back to the dissociated single cells, we mapped the homeobox transcription factor gene *cdx4* and several *hox* genes to assign whether a cluster likely represented anterior or posterior cells within the embryo (**Fig. 2D, Fig. S3**). *cdx4* at 5 somite stages broadly demarcates in the posterior half of the developing zebrafish embryo (**Fig. 2D**). *cdx4*-positive and thus posterior clusters composed of cells expressing marker genes for endothelial and hematopoietic precursors (*fli1a, lmo2, znfl2a*) (Hogan et al., 2006; Lawson and Weinstein, 2002; Zhu et al., 2005), and for the pronephros (*pax2a, pax8, lbx2*) (Naylor et al., 2016; Perens et al., 2016; Pfeffer et al., 1998) (**Fig. 2E**). The overlapping expression of hemangioblast and kidney markers possibly indicates that these two clusters represent a mixed multi-lineage progenitor pool at the end of gastrulation.

Notably, among all the prominent LPM genes we detected *hand2* expression across several clusters. In addition to clusters encompassing the expected cardiopharyngeal precursors, *hand2* transcripts were abundant in four clusters we accordingly named *hand2*-high_1-4 (**Fig. 2C,E**). Further, expression of *hand2* fell into both anterior and posterior expression domains, correlating with its native expression pattern in zebrafish (Yelon et al., 2000) (**Fig. 2C,E**). As anticipated, *hand2* transcripts coincided with the expression of cardiac genes (**Fig. 2E**). However, the anterior *hand2*-high_3 and *hand2*-high_4 clusters showed no detectable expression of cardiac markers including *hey2, tbx20, nkx2.5*, and *nkx2.7*, suggesting that they encompass cells with another fate potential. We also found posterior *hand2*-positive cells that appeared distinct from endothelial, hematopoietic, and kidney progenitors (*hand2*-high_1 and *hand2*-high_2) (**Fig. 2E**). Taken together, our scRNA-seq captured *hand2* expression as a central feature of several LPM progenitor clusters.

We next aimed to determine what genes are co-expressed with *hand2*. Our analysis revealed that several genes were enriched and individually even among the cluster-defining genes in one or more of the *hand2_high* clusters, including *sfrp5, foxh1, gata5, gata6*, and *meis3* (**Fig. 2F-K, Fig. S4**). In fluorescent mRNA ISH (RNAscope) and colorimetric mRNA ISH on whole-mount tailbud and early somitogenesis staged embryos, we observed that the endogenous expression pattern of *meis3* expression overlaps with the *hand2* domain in the anterior and posterior LPM, among with *meis3* expression in other non-LPM domains (**Fig. 2G,I**). Moreover, *foxh1* expression has previously been described in the LPM during somitogenesis stages (Pogoda et al., 2000; Slagle et al., 2011), yet the exact domain within the LPM has remained unclear. Akin to *meis3*, our fluorescent ISH confirmed that *foxh1* is expressed in the lateral-most posterior LPM territory overlapping with *hand2* expression (**Fig. 2H**). Notably, the in part redundant transcription factor genes *gata4, gata5*, and *gata6* that play key roles in cardiac and endoderm development (Tremblay et al., 2018), are also expressed in a domain lateral to the forming heart field (Jiang et al., 1999; Reiter et al., 1999). The cells in this domain take on a spread-out, mesh-like pattern over the yolk during somitogenesis (**Fig. 2J**). Similarly, we found that *sfrp5* demarcates lateral-most LPM cells that form a spread-out expression domain along the body axis over the yolk and yolk extension (**Fig. 2K**). This mesh-like pattern can also be recognized within the endogenous expression of *hand2* within the ALPM (Yelon et al., 2000). Taken together, our data documents a collection of genes co-expressed with *hand2* in the lateral-most LPM domain of a yet-unknown fate.

### *hand2* expression identifies mesothelial progenitors

Our analysis indicated that expression of *hand2* and several associated genes demarcate a lateral-most LPM domain that is distinct from the cardiopharyngeal, blood, vasculature, and kidney progenitors at the end of gastrulation. Based on the previous association of Hand1/2 expression with differentiated mesenchymal structures in mouse and chick (Ahn et al., 2002; Charité et al., 2000; Fernandez-Teran et al., 2000; Firulli et al., 1998; Perens et al., 2016; Prummel et al., 2019), we hypothesized that the *hand2*-expressing LPM in zebrafish, in particular the posterior lateral-most stripe, forms the mesothelium. We turned to the transgenic line *hand2:EGFP* based on a BAC encompassing the zebrafish *hand2* locus that faithfully recapitulates endogenous *hand2* expression (Yin et al., 2010). Using time-lapse SPIM imaging and panoramic projections, we captured the dynamics of *hand2:EGFP* reporter activity during segmentation stages, demarcating the lateral-most *drl*-expressing LPM (**Fig. 3A,B**) (Perens et al., 2016; Prummel et al., 2019). Notably, and in contrast to the more medial LPM stripes that progressively migrated to the midline, a subset of the *hand2*:EGFP-expressing cell population sprawled out laterally over the yolk as a single-cell layer (**Fig. 3C**).

**Figure 3:**
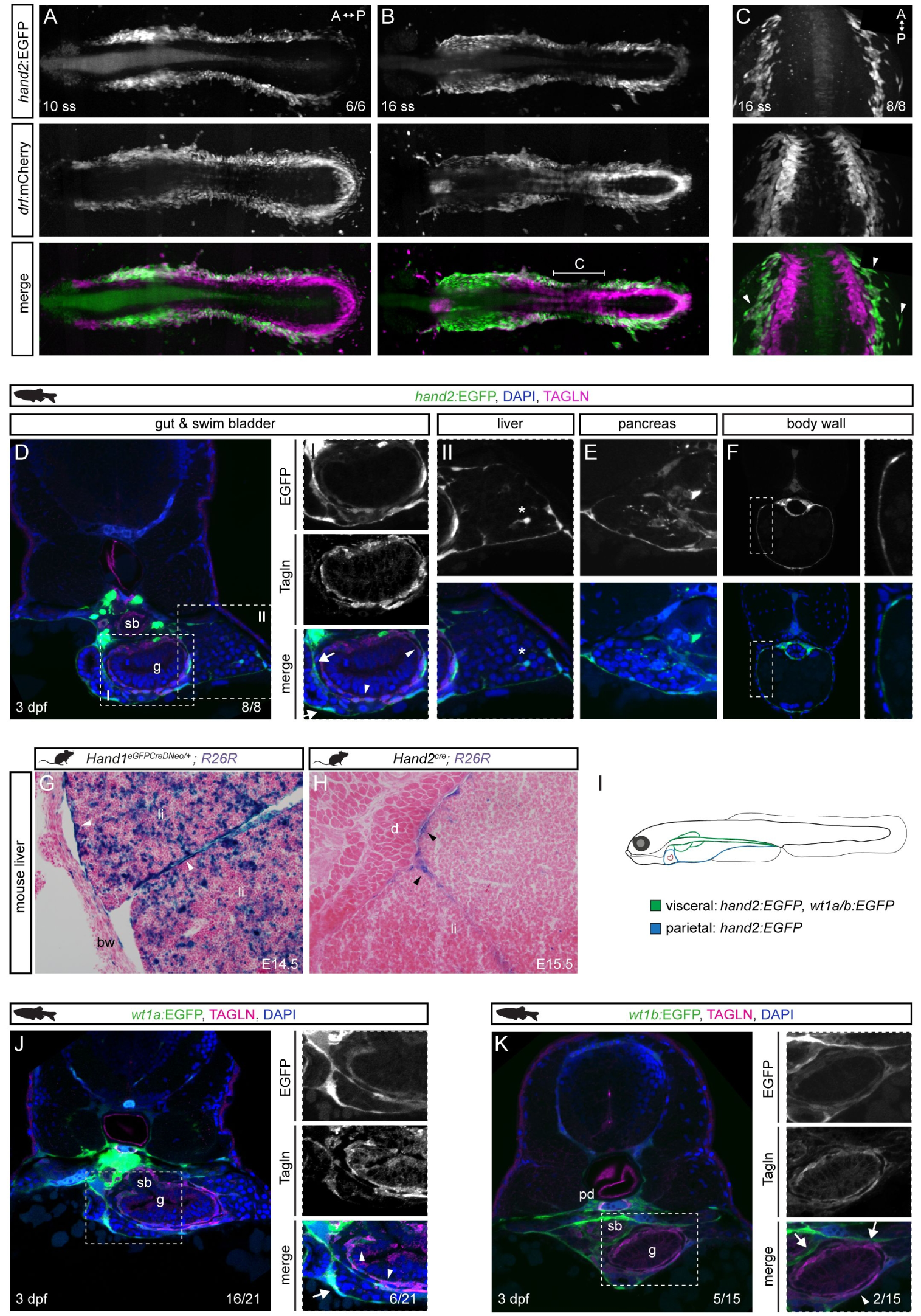
*hand2* defines mesothelial precursors in the LPM. **(A,B)** SPIM projections of *hand2:EGFP;drl:mCherry* embryos at 10 ss (**A**) and 16 ss (**B**). **(C)** Confocal imaging of 16 ss *hand2:EGFP;drl:mCherry* embryo showing *hand2:EGFP*-expressing cell populations comprising lateral-most LPM. Arrowheads label single-cell layer laterally migrating over yolk. Region as annotated in (**B**). **(D-F)** Transverse sections of 3 dpf *hand2:EGFP* embryos. **(D)** Boxed regions show double-positive Tagln staining (arrowheads) and Tagln-negative staining in visceral peritoneum around (I) gut, pancreatic/hepatic ducts (arrows), and (II) liver. Asterisk depicts potential hepatic stellate cell. **(E)** *hand2*:EGFP expression in visceral peritoneum surrounding the pancreas. **(F)** Transverse section of *hand2:*EGFP labeling of parietal peritoneum. (**G,H**) Abdominal transverse sections of E14.5 *Hand1*^*EGFPCreΔNeo/+*^;*R26R* and E15.5 *Hand2*^*Cre*^;*R26R* mouse embryos, lineage-labeled cells marked by β-galactosidase staining (blue). In both groups, lineage labeling appears in the visceral peritoneum of the liver (arrowheads). (**I**) Illustrated mesothelial layers in 3 dpf zebrafish embryo. **(J,K)** *wt1a:*EGFP (**J**) and *wt1b:EGFP* (**K**) expression in pronephric ducts and visceral peritoneum surrounding the gut, swim bladder, and liver. Boxed regions show double-positive Tagln staining (arrowheads) and Tagln-negative staining (arrows). Body wall (bw), diaphragm (d), gut (g), liver (li), pronephric duct (pd), swim bladder (sb). Nuclei in blue (DAPI). Scale bar (**D,F,J,K**) 50 μm, (**E, boxed regions D,F,J,K**) 25 μm.

In transverse sections of *hand2:EGFP-*transgenic embryos at 3 dpf, we observed the previously described *hand2* reporter expression in Tagln-positive, LPM-derived intestinal smooth muscle cells that layer around the endodermal gut tube (Gays et al., 2017) (**Fig. 3D**). In higher magnification analysis of transverse sections, we also observed Tagln-negative *hand2:*EGFP-expressing cells surrounding the gut at 3 dpf, reminiscent of coelomic epithelial cells (**Fig. 3D**). At 3 dpf, the *hand2* reporter-expressing layers had also wrapped around other endoderm-derived organs, including the liver and the pancreas (**Fig. 3D,E**). In addition, in transverse sections capturing the yolk at 3 dpf, we detected *hand2*:EGFP expression underneath the skin in the body wall, the prospective parietal peritoneum (**Fig. 3F**). Extending previous observations (Gays et al., 2017; Mao et al., 2015; Perens et al., 2016; Yin et al., 2010), these data suggest that *hand2:EGFP* reporter expression at 3 dpf in zebrafish delineates the visceral and parietal peritoneal membranes.

To extend our interpretation of zebrafish *hand2* as marker for the emerging mesothelium, we next turned to the mouse to uncover any mesothelial lineage contribution of cells expressing either of the two partially redundant murine *Hand* genes *Hand1* and *Hand2*. Crossing either the *Hand1*^*EGFPCreΔNeo/+*^ (Barnes et al., 2010) or *Hand2*^*Cre*^ (Ruest et al., 2003) transgenic strains into the *R26R loxP* reporter strain (Soriano, 1999) resulted in lineage labeling along the mesothelium lining of the liver lobes at E14.5 and E15.5, respectively (**Fig. 3G,H**). We found lineage labeling to be more apparent and widespread using the *Hand1*^*EGFPCreΔNeo/+*^ strain, with strong staining in most of the mesothelium and in a subset of cells within the liver of currently unknown identity (**Fig. 3G**). Labeling was also present in some cells lining sinusoids, though more sporadic (data not shown). Lineage analysis of *Hand2*^*Cre*^-descendant cells also revealed lineage labeling in the mesothelium (**Fig. 3H**), though staining was more restricted and less robust than that observed in *Hand1* daughter cells. These differences could reflect differences between the two Cre drivers (knockin vs transgenic) or a larger proportion of *Hand1*-expressing cells contributing to the liver mesothelium.

To further investigate a possible conserved link of *Hand* gene expression to mesothelium formation in mouse, we mined recent scRNA-seq and scATAC-seq data of early mouse embryos (Pijuan-Sala et al., 2020) (**Fig. S5**). This data harbored *Hand2-* expressing cell clusters that were previously assigned as “mesenchymal” and that share co-expression of transcription factor genes we found in our zebrafish LPM dataset as associated with mesothelial progenitors, including *Meis3, Gata5*, and *Gata6* as well as the previously LPM-associated genes *Gata4, FoxF1*, and *Hand1* (Prummel et al., 2020) (**Fig. 2, Fig. S5**). Together, these observations indicate that also in mammals *Hand* gene expression is a conserved feature of the mesothelial progenitors.

To further confirm and define the *hand2*-expressing mesothelia, we turned to Wt1 expression that characterizes the developing visceral mesothelium in mammals (Chen et al., 2014; Parenti et al., 2013; Walker et al., 1994). Expression of the zebrafish *Wt1* paralogs *wt1a* and *wt1b* becomes detectable at 6-8 ss as previously analysed in kidney and epicardium development (Bollig et al., 2009; Endlich et al., 2014; Peralta et al., 2014; Perner et al., 2016). During somitogenesis, we found *wt1a:EGFP* and *wt1b:EGFP* activity lateral of the differentiating kidney structures within the *drl:mCherry-*expressing LPM corresponding to the *hand2*-positive territory (**Fig. S6A,B**). In transverse sections of *wt1a-* and *wt1b:EGFP*-expressing embryos, we detected EGFP signal in the coelomic epithelium surrounding the gut, liver, and pancreas in addition to the previously reported labeling of the glomerulus and pronephric tubules (**Fig. 3J-K**) (Bollig et al., 2009; Endlich et al., 2014; Peralta et al., 2014; Perner et al., 2016). Further, we observed *wt1a/b* reporter-expressing Tagln-positive smooth muscle cells around the gut and hepatic and pancreatic ducts (**Fig. 3J-K**). Genetic lineage tracing using *wt1a*- and *wt1b:creERT2* from the onset of transgene expression at 6-8 ss robustly marked the visceral peritoneum around the gut, liver, and pancreas (**Fig. S6C-E**). Notably, at 3 dpf, we neither detected any EGFP-labeling of the visceral peritoneum around the more posterior gut, nor did we detect any EGFP-expressing parietal peritoneum cells around the yolk, indicating these cells do not express *wt1* genes in zebrafish at our analyzed time points (**Fig. S6F-G**). Extending previous *wt1a*-based lineage tracing (Sánchez-Iranzo et al., 2018a), we observed that also *wt1b*-expressing cells contribute to the cardiac mesothelial layers in the dorsal and ventral pericardium, and in the pro-epicardial clusters (**Fig. S6H,I)**. These observations are consistent with LPM lineage tracing using *drl:creERT2* to ventral and dorsal pericardium and to proepicardium (**Fig. 1F, Fig. S1D-I**). We conclude that, akin to the mammalian mesothelium, expression of both *Wt1* orthologs is also a feature of developing mesothelia in zebrafish. Nonetheless, in addition to the absence of *wt1a/b* expression in the parietal peritoneum lineage, *hand2:EGFP* presents an earlier and more complete marker of all developing mesothelial membranes in zebrafish.

### Primordial germ cells associate with mesothelial progenitors

We noted the chemokine-encoding gene *cxcl12* significantly co-expressed with *hand2* in our scRNA-seq data set (**Fig. 2E, Fig. 4A**). Cxcl12a provides directional cues for the migration of several cell types (Lewellis and Knaut, 2012), including the guidance of primordial germ cells (PGCs) (Grimaldi and Raz, 2020). Through complex migration paths, the PGCs reach their final destination in the region where the gonad develops, dorsally and medially bordered by the developing pronephros, while the identity of the ventrally located tissue has not been precisely defined (Doitsidou et al., 2002; Richardson and Lehmann, 2010; Stebler et al., 2004). We asked whether the *drl*- and *hand2*-expressing mesothelium interacts with the migrating PGCs and if PGCs are possibly already associated with the developing LPM during gastrulation. In time-lapse imaging of transgenic zebrafish embryos expressing EGFP in the LPM (*drl:EGFP* transgene) and the farnesylated mCherry in their PGCs (*kop:mCherry-f’-nos3’UTR* transgene) (Blaser et al., 2005), we observed that the migrating PGCs associate with the *drl* reporter-expressing mesendoderm already during gastrulation (**Fig. 4B,C, Movie 1**). The *drl:EGFP*-expressing progenitor field is gradually specifying into the LPM at the beginning of somitogenesis (Prummel et al., 2019) and the PGC clusters continued to associate with, and migrate within, the forming LPM **(Fig. 4B,C**). By 24 hpf, the PGCs reach the region where the gonads will form (Raz, 2003) and we speculated the *hand2*-positive mesothelium-primed LPM serves as the final destination for the PGCs. Imaging *mCherry-f’-nos3’UTR*;*hand2:EGFP* embryos documented how the PGCs migrated within the *hand2:EGFP*-expressing LPM, and the PGCs eventually arrive at a domain within the *hand2*-expressing forming mesothelium (**Fig. 4D**). Antibody staining for the kidney transcription factor Pax2a confirmed that the PGCs end up ventral of the pronephros within the *hand2*-positive LPM (**Fig. 4,EF**), with the developing gut located medially between the PGC clusters (Paksa et al., 2016). These observations further support our conclusion that the PGCs migrate in close association with the forming coelomic epithelium, as marked by *hand2* expression, already during early development.

**Figure 4:**
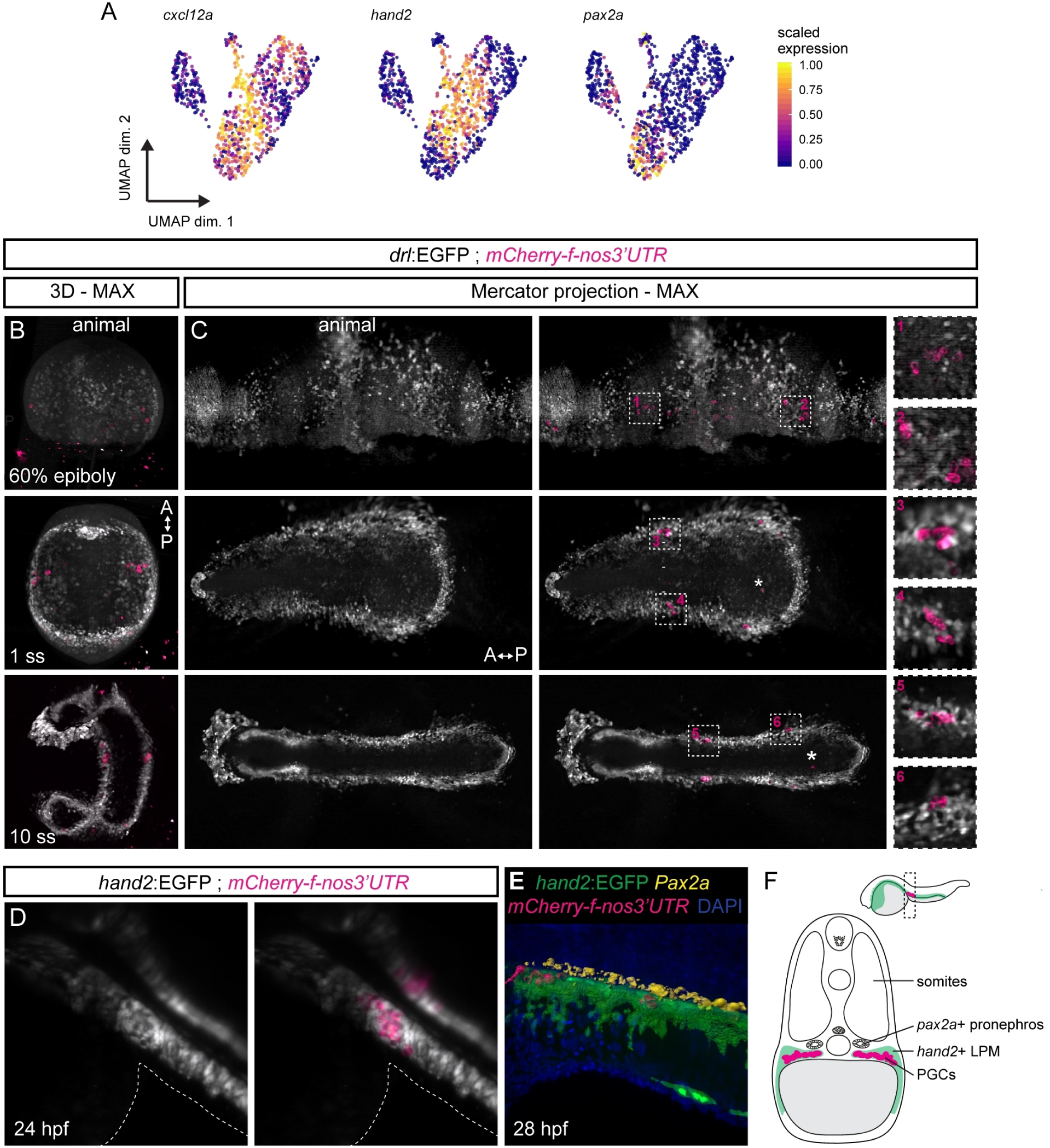
Primordial germ cells migrate with, and home within, the *hand2*-positive LPM. **(A)** UMAP plots of *cxcl12a, hand2*, and *pax2a*. Cells are colored by scaled expression values using lower and upper 1%-quantiles as boundaries. **(B,C)** Snapshots of time-lapse movie (**Movie 1**) showing *drl:EGFP* with primordial germ cell (PGC) marker *mCherry-f’-nos3’UTR*, with 3D renderings at 60% epiboly, 1 ss, and 10 ss (**B**) and maximum intensity Mercator projections for the same time points (**C**). The PGCs migrate from four different clusters during gastrulation into two bilateral clusters during somitogenesis (boxed regions). Note the sparse (lost) PGCs located within the endoderm (asterisks). Animal side is for 60% epiboly to the top, anterior for 1 ss and 10 ss in (**B**) to the top and in (**C**) to the left. **(D)** Imaging of *hand2:EGFP* with *mCherry-f’-nos3’UTR* showing a dorsal view at 24 hpf: the PGC clusters fall completely within the *hand2:EGFP* domain. The outlines of the yolk extension are highlighted. **(E)** Segmented 3D rendering of a *hand2:EGFP; mCherry-f’-nos3’UTR* double-positive embryo stained with anti-Pax2a (yellow) and DAPI (blue) at 28 hpf. **(F)** A schematic transverse section of a 24 hpf embryo showing two bilateral PGC clusters (magenta) lateral of the developing gut, ventral-lateral of the *pax2a*-positive developing pronephros (yellow) within the *cxcl12a*-high *hand2*-positive LPM/coelomic epithelium (green).

### *hand2*-expressing LPM forms the visceral and parietal mesothelium

To clarify their mesothelial identity and to resolve the dynamics of *hand2:EGFP*-expressing mesothelial progenitor cells in the zebrafish embryo, we performed long-term and *in toto* light sheet imaging of developing *hand2:EGFP;drl:mCherry* double-transgenic embryos. Applying a multi-sample imaging and processing workflow (Daetwyler et al., 2019), we captured the dynamics of the *hand2:EGFP* expression over the course of embryonic development from 18 to 82 hpf *in toto* (n = 6 embryos). At imaging onset, the *hand2:*EGFP-expressing cells had initiated their lateral migration over the yolk, while expression of *drl:mCherry* refined to cardiovascular and blood lineages (**Fig. 5A, Movie 2**) (Mosimann et al., 2015).

**Figure 5:**
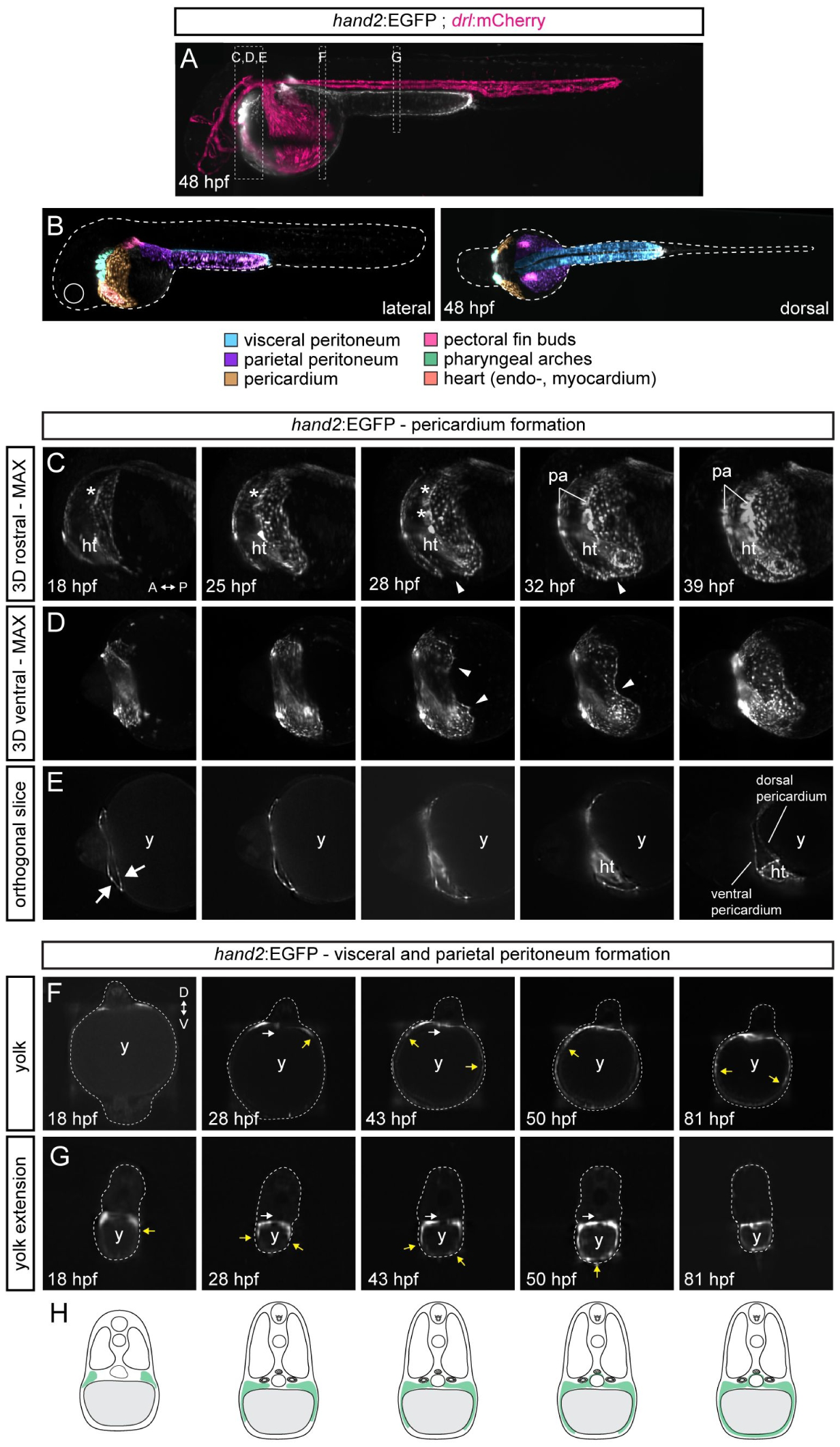
Formation of the *hand2*-positive mesothelium. Multiday, multi-angle SPIM of *hand2:EGFP* zebrafish embryos from 18-82 hpf (n = 6). **(A)** Maximum intensity projection (MAX) of embryo expressing the LPM-marking *drl:mCherry* (24 hpf onwards restricted to cardiovascular lineages) (magenta) and *hand2:EGFP* (greyscale) at 48 hpf. Boxes depict selected regions in C-G. **(B)** *hand2*-positive populations within 48 hpf embryo, lateral view left and dorsal view right. **(C-E)** 3D-rendered *hand2:EGFP* embryo, focused on pericardium formation. (**C**) Rostral view, visualizing formation of pharyngeal arches (asterisks) and primary heart tube. (**D**) Ventral view, illustrating left and right flanks of the forming pericardium meeting at the midline (32 hpf, arrowheads). (**E**) Single plane, highlighting how the pericardial cavity forms within anterior LPM (arrows). *hand2* labels coelomic epithelium, contributing to ventral and dorsal pericardium. **(F,G)** Single-plane cross-sections showing the migration of *hand2*-positive cells over yolk (**G**) and yolk extension (**H**), forming parietal peritoneum. Yellow arrows point out dorsal-ventral directed migration path. White arrows indicate inwards-migrating EGFP-expressing cells, contributing to coelomic epithelium maturing into visceral peritoneum. **(H)** Schematics of how *hand2*-expressing cells laterally migrate over the yolk forming parietal peritoneum, and migrating medially to wrap around endodermal-derived organs forming the visceral peritoneum. Red blood cells (rbc), heart tube (ht), pharyngeal arches (pa), yolk (y).

In the resulting time lapse imaging, we captured the *hand2:EGFP*-positive LPM as a dynamic cell population with distinct migration behaviors in different regions of the developing embryo. First, we observed the *hand2:EGFP*-expressing cells contributing to the previously described processes of pharyngeal arch formation, pectoral fin bud outgrowth, and primary heart tube extension (**Fig. 5B, Movie 2,3**). Additionally, our light sheet imaging registered that two seemingly separated populations of *hand2*:EGFP-expressing cells migrate ventro-laterally, crawling over the yolk and yolk extension, and continue to form the parietal layer of the cardiac cavity (pericardium) and abdominal cavity (parietal peritoneum), respectively (**Fig. 5C-G**). Already at 18 hpf, we observed a dorso-ventral split in the *hand2*-expressing progenitor field positioned lateral of the cardiopharyngeal progenitors, forming the cardiac cavity (**Fig. 5E**). The dorsal and ventral pericardium surround the forming heart tube and maintain the pericardial fluid that supports the beating, moving heart (Peralta et al., 2013). The cells that will form the dorsal pericardium migrated over the yolk as a connected cell field with a clearly visible leading edge and were already interacting with the forming primary heart tube (**Fig. 3B, 5D, Movie 2,3**). We observed that the cells forming the more posterior-located parietal peritoneum are split from the forming pericardium by the circulation valley forming over the anterior yolk, where later the common cardinal vein (Duct of Cuvier) migrates into to ultimately enclose the circulating blood (Hamm et al., 2016) (as visualized by *drl:*mCherry, **Movie 1, Fig. S7**). In addition, we documented how the *hand2*:EGFP-cells forming the presumptive parietal mesothelium reach the ventral midline earlier around the caudal yolk extension than around the main yolk, possibly due to the shorter migration distance over the yolk extension’s smaller diameter (**Fig. 5F-H**). Simultaneously, the more medially located *hand2:EGFP-*expressing cells migrated towards the midline (**Fig. 5F-H**), corresponding to the formation of visceral peritoneum including intestinal smooth muscles (Gays et al., 2017; Yin et al., 2010). Taken together, these imaging data further support the notion that *hand2:EGFP*-expressing LPM progenitors in zebrafish form the visceral, parietal, and pericardiac mesothelium.

### *hand2* mutants fail to form their mesothelia and to close the ventral body wall

The prominent expression of *hand2* in mesothelial progenitors of the LPM prompted us to investigate whether the loss of *hand2* has an impact on the formation of the mesothelial layers. Phenotype analyses of mutant mice and zebrafish have documented that Hand2 (partially redundantly with Hand1 in mice) contributes to the development of the heart, forelimbs/pectoral fins, and smooth muscle lineages (Galli et al., 2010; Laurent et al., 2017; Santoro et al., 2009; Srivastava et al., 1997; Yelon and Stainier, 2005; Yelon et al., 2000). Additionally, zebrafish *hand2* mutants feature disorganized migration of the intestinal smooth muscle progenitors towards the gut and overall gut mislooping due to disorganized extracellular matrix remodeling (Gays et al., 2017; Yin et al., 2010).

We revisited the previously established zebrafish mutant *han*^*S6*^ that harbors a presumptive *hand2* null allele (Yelon et al., 2000). In addition to the well-described cardiac and pectoral fin defects, we observed that *han*^*S6*^-homozygous embryos displayed ventral defects of their yolk extensions and subsequent deterioration of the embryo due to yolk herniation at 3 dpf (**Fig. 6A-D**). Moreover, we noticed selective blistering and shriveling of the ventral fin fold. At the cellular level, in contrast to the connected epithelial layer forming mesothelia in wildtype embryos, we observed that LPM-derived cells (marked by genetic lineage tracing using *drl:creERT2*) are sparse in the abdominal cavity in *han*^*S6*^-homozygous embryos (**Fig. 6E**). The sparse distribution of LPM-derived cells could potentially result from mosaic labeling from incomplete *loxP*-reporter recombination following *drl*-driven CreERT2 activation, as common in Cre/*loxP*-based experiments (Carney and Mosimann, 2018). However, *han*^*S6*^-homozygous embryos displayed similar migration issues when we directly visualized for the mesothelial cells with the *wt1a:EGFP* and *hand2:EGFP* transgenic reporters, respectively (**Fig. 6F,G**). We therefore conclude that loss of *hand2* perturbs mesothelial progenitor migration, in line with previous observations indicating that coelomic epithelial progenitors migrated to the midline in *hand2* mutants but failed to properly wrap around the gut (Yin et al., 2010). Of note, we only observed few LPM lineage-labeled cells and *hand2:*EGFP-expressing cells lining the body wall, respectively, indicating that also the ventrally migration of the parietal peritoneum progenitors is perturbed in *hand2* mutants (**Fig. 6E,G**). Consequently, *han*^*S6*^-homozygous embryos also fail to form a proper body wall, likely resulting in herniation of the yolk along the anterior-posterior axis over time. These observations in zebrafish connect *hand2* expression and function in dedicated LPM progenitor cells to the proper execution of mesothelial membrane formation.

**Figure 6:**
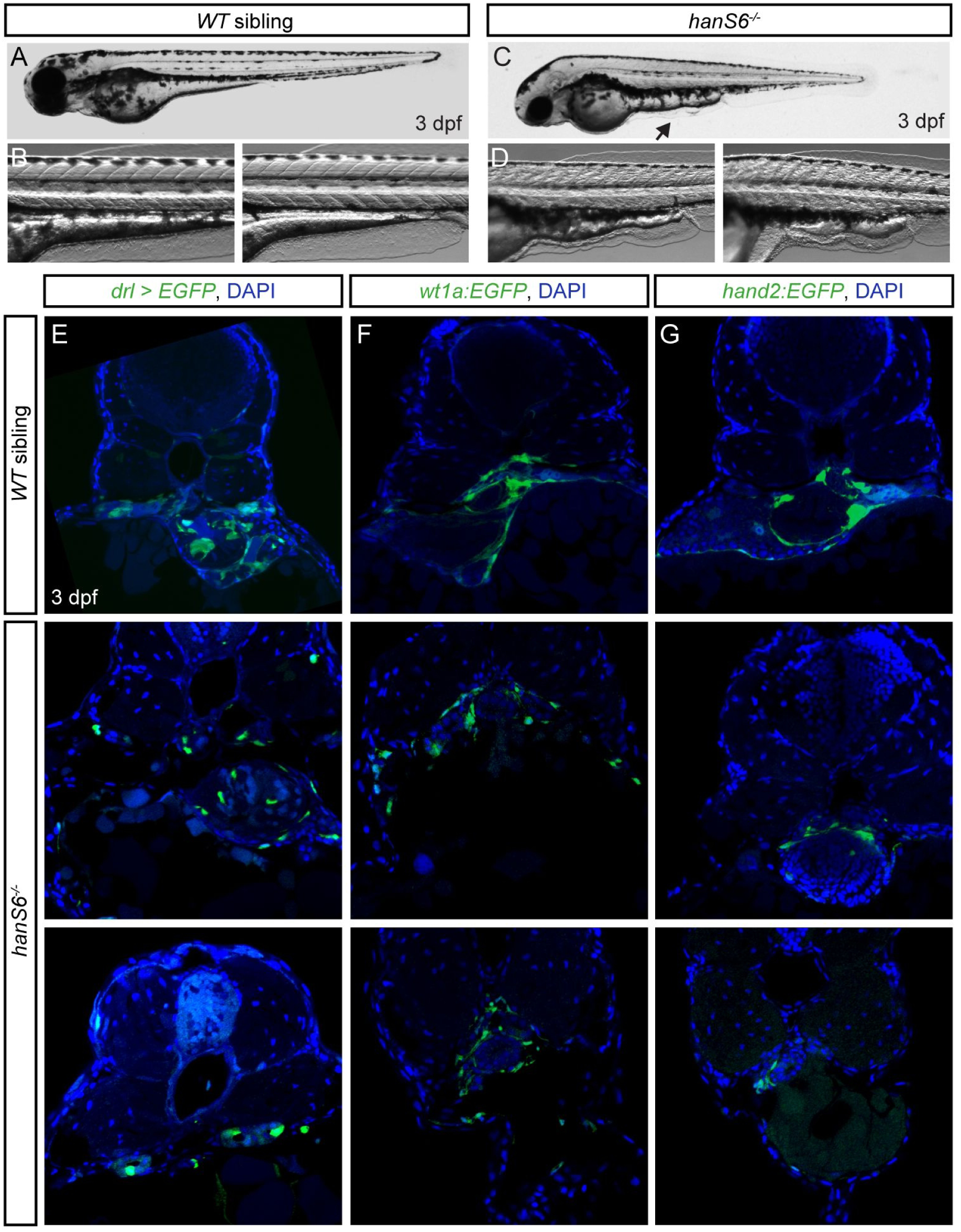
Loss of *hand2* causes peritoneum defects. **(A-D)** Lateral views of a phenotypically wild-type (**A**) sibling and representative *han*^*S6*^ mutant (**C**) at 3 dpf. Note the abnormal phenotype of the yolk extension in the mutant embryo (arrow), what is in additionally highlighted in the zoom-ins of two representative pictures of wild-type siblings (**B**) and *han*^*S6*^ mutants (**D**). **(E-G)** *han*^*S6*^ in the background of *drl:creERT2;hsp70l:Switch* (**E**), *wt1a:EGFP* (**F**), and *hand2:EGFP* (**G**). Visceral peritoneum is disorganized around the internal organs and the body wall and the associated parietal peritoneum are malformed (**G**).

### Mesothelioma activates early LPM transcription factors including Hand2

Upon exposure to asbestos fibers, mesothelial cells within the visceral and parietal pleura can transform into mesothelioma tumors (Carbone et al., 2019). As the cell(s) of origin have been hypothesized to reside within the mesothelium, we sought to determine whether mesothelioma tumors feature a transcriptional signature akin to the mesothelial progenitors we found in zebrafish (**Fig. 2**).

First, we revisited our transcriptomics data set obtained from mouse mesothelioma triggered by repeated exposure to crocidolite fibers (blue asbestos) (Rehrauer et al., 2018) (**Fig. 7A**). As previously established, genes found upregulated in human mesothelioma development (including *Msln* and *Wt1*) or downregulated (including tumor suppressors *Nf2* and *Bap1*) behaved analogously in this mouse model, both in pre-neoplastic lesions and in fully formed tumors when compared to healthy adult mesothelium (**Fig. 7B**) (Bianchi et al., 1995; Bott et al., 2011; Bueno et al., 2016; Hmeljak et al., 2018; Rehrauer et al., 2018; Sekido et al., 1995). Early LPM lineage markers, including *Scl/Tal1, Etv2*, and *Pax2*, were not detected with RNA-seq in the adult mouse mesothelium samples. Notably, the sham-treated mesothelial tissue also expressed no or low levels of transcripts for LPM-associated genes including *Hand2, Gata4/5/6, Meis3, FoxF1, Mixl1*, and *Lmo2* (**Fig. 7B**). In contrast, the transition to pre-neoplastic lesions and to mesothelioma after crocidolite exposure was accompanied by the upregulation of mouse orthologs of several genes associated with mesothelium progenitors in zebrafish: expression of *Hand2, Wt1, Gata4/5/6*, and *Meis3* increased in crocidolite-exposed mesothelium, with a particularly striking upregulation in fully formed tumors (**Fig. 7B**). Also *FoxF1* was upregulated in crocidolite-induced tumors, a gene which has been associated with the developing splanchnic mesoderm in lamprey, chicken, and mouse, but not in zebrafish (Funayama et al., 1999; Mahlapuu et al., 2001; Onimaru et al., 2011; Ormestad et al., 2004) (**Fig. 7B**). Following immunohistochemistry for Hand2, we detected specific and prominent nuclear Hand2 immunoreactivity in crocidolite-induced mesothelioma tumor sections (**Fig. 7C**). Altogether, these observations suggest that the transformative events leading to mesothelioma in a crocidolite-based mouse model are accompanied by the upregulation (or re-initiation) of early LPM genes we found associated with mesothelial progenitors in development (**Fig. 2**).

**Figure 7:**
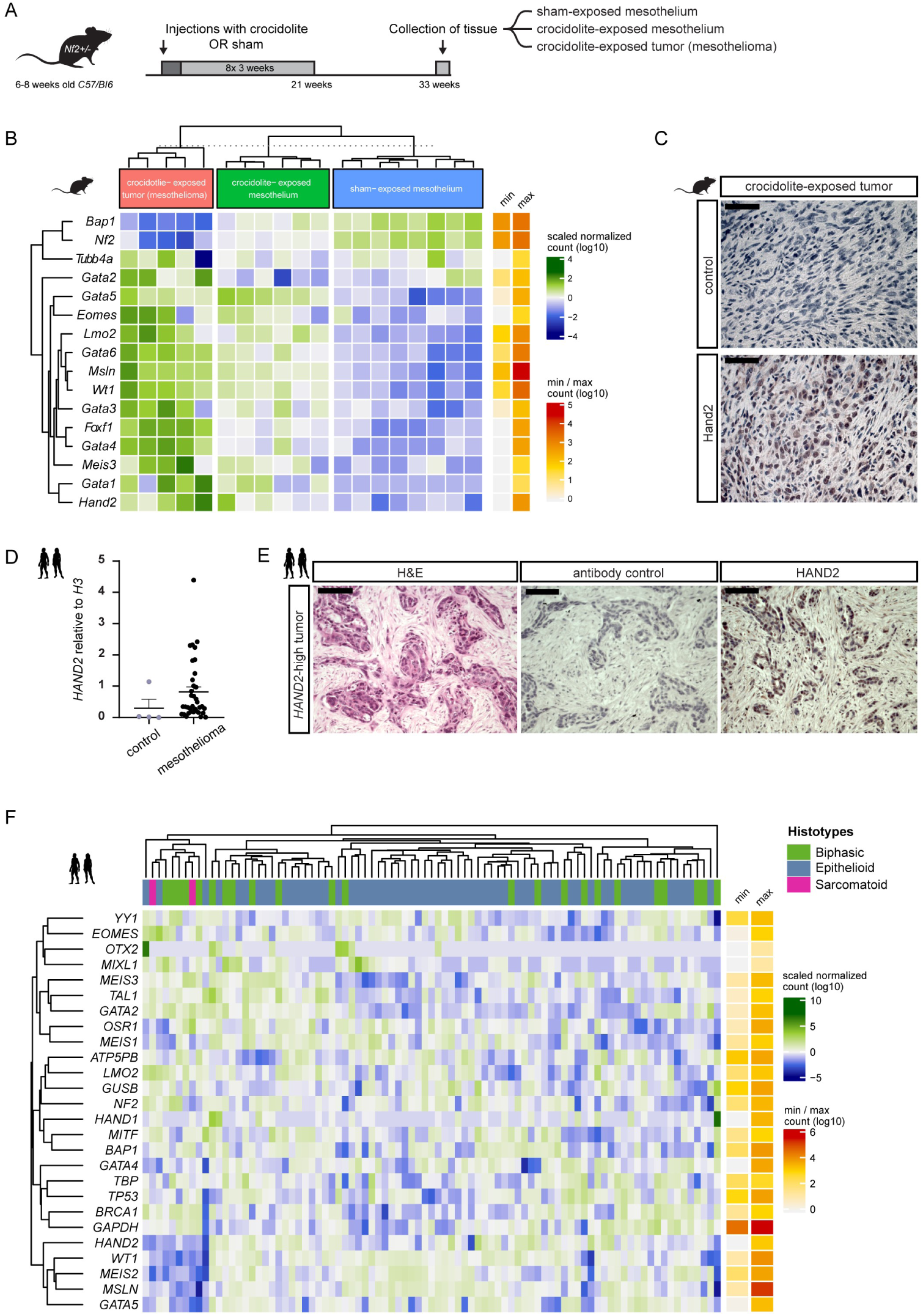
Mesothelioma reactivates an early LPM program. **(A)** Modeling loss of mesothelial homeostasis during mesothelioma tumorigenesis: 6-8 weeks-old *Nf2*+/-*C57Bl/6J* mice exposed to crocidolite intra-peritoneally (400 µg) every 3 weeks with 8 total treatments, parallel to controls. 33 weeks after initial exposure, tissue was collected. (**B**) Heatmap of LPM-associated genes, either up-(*Msln, Wt1*) or downregulated (*Nf2, Bap1)* mesothelioma genes, and negative control (*Tubb4a*). Bins colored by row-scaled log2-normalized counts; columns (samples) split by treatment group; rows and columns ordered by hierarchical clustering (scaled expression values). Right side row indicates raw count range. (**C)** Immunoreactivity of Hand2 protein in mouse mesothelioma. (**D**) Relative expression of *HAND2* by qPCR in normal pleura (n = 4) and human malignant mesothelioma (n = 36); trend (p = 0.0596) towards increased *HAND2* levels in tumors (Mann-Whitney test). Mean ± SE. (**E)** Immunoreactivity of HAND2 protein in *HAND2*-high human mesothelioma. (**F**) Heatmap representing unsupervised hierarchical clustering of human mesothelioma from TCGA RNA-seq (n = 87), depicting expression of mesothelial progenitor-associated LPM genes, mesothelioma-associated genes (*WT1, MSLN, BAP1, NF2, TP53*), unrelated control (*YY1, OTX2, BRCA1*) and housekeeping genes (*ATP5PB, GUSB, TBP, GAPDH*). Columns represent histotypes, rows represent log10-transformed, batch-normalized mRNA expression levels of mesothelioma samples. Scale bars **(C)** 50, **(D)** 100 µm.

We next sought to corroborate whether human mesothelioma also feature *HAND2* expression. Analyzing expression levels of *HAND2* using qPCR in human pleural mesothelioma (n = 36) compared to healthy pleura (n = 4), we observed that individual tumors showed heterogeneous yet consistent upregulation of *HAND2*, with individual tumors expressing high levels of *HAND2* mRNA (p = 0.0596) (**Fig. 7D)**. We further detected nuclear HAND2 immunoreactivity in an epithelioid mesothelioma sample with high *HAND2* mRNA levels (**Fig. 7E**). To further establish if expression of HAND2 and other mesothelial progenitor-associated LPM genes are re-activated in mesothelioma, we mined the mesothelioma-associated transcriptome data deposited in the The Cancer Genome Atlas (TCGA) as generated by the TCGA Research Network (https://www.cancer.gov/tcga). We compared gene expression for *HAND2* and other mesothelial progenitor genes with a collection of mesothelioma-associated genes, and several tissue-specific and ubiquitous house-keeping genes. We found that increased or re-activated *HAND2* expression was not an obvious signature across the analyzed mesothelioma samples, but rather a feature of a subset of tumors across the epithelioid, sarcomatoid, and biphasic classifications (**Fig. 7F**). *HAND2* expression coincided with differential expression of the previously mesothelium- and mesothelioma-associated genes *MSLN* and *WT1*, as well as with *GATA5* and *MEIS2* (**Fig. 7F**). Notably, while we found in zebrafish mesothelial progenitors *meis3* as the most-prominent *MEIS* family gene co-expressed with *hand2* (**Fig. 2**), its redundant paralogs *meis2a* and *meis2b* were also enriched in *hand2*-expressing zebrafish progenitors (**Fig. S4**) (Schulte and Geerts, 2019). This *HAND2*-expressing group of mesothelioma tumors did not cluster notably with tumors featuring differential expression of the common mesothelioma-associated tumor suppressor genes *BAP1* and *NF2* (**Fig. 7F**). Taken together, these observations provide first indications that mesothelioma tumors can upregulate or reactivate several developmental factors associated with an mesothelium progenitor state, in particular Hand2 that in zebrafish provides the earliest mesothelium progenitor marker. We postulate that, following malignant transformation, mesothelioma tumors acquire features of a developmental progenitor program deployed by the early LPM to specify mesothelium progenitors, endowing transformed cells with unique migratory and lineage properties.

## Discussion

Despite their various contributions to development and homeostasis, the early embryonic origins of mesothelia and their genetic control have remained uncertain. Uniquely positioned within the developing embryo to cover inner organs and the forming body cavities, our results uncover that mesothelial progenitor cells emerge among the earliest distinguishable LPM cell fates following *hand2* expression.

Our finding that expression of the transcription factor Hand2 demarcates the merging mesothelial progenitors within the post-gastrulation LPM provides new means to study the mesothelium in development and disease. *Hand* gene function centrally contributes to cardiac, limb, and pharyngeal arch development across several models (Barnes et al., 2011; Firulli et al., 1998; Han and Olson, 2005; Shin et al., 2009; Yelon et al., 2000). Our results indicate that *Hand* genes also act as conserved regulators of mesothelial fates. Gene expression studies have defined *hand2* as a marker for the forming body wall in tilapia and medaka, and have found *Hand1/2* in the flank of the developing mouse embryo (Srivastava et al., 1995; Srivastava et al., 1997; Tanaka, 2011; Tanaka et al., 2015). Complementing studies in mice have demonstrated that *Hand1*-expressing progenitors contribute to cell layers of the gut including smooth muscles (Barnes et al., 2010; Srivastava et al., 1995; Srivastava et al., 1997). A possible ancient association of *Hand* genes with the LPM and thus mesothelium progenitors followed the expression of amphioxus *AmpiHand* and lamprey *LjHandA* within their presumptive LPM (Onimaru et al., 2011). While zebrafish have seemingly lost their *Hand1* ortholog (Howe et al., 2013), our data suggest an early role for Hand2 and Hand factors in general in the differentiation of mesothelial progenitors from the LPM (**Fig. 3**).

Several fates emerging from the LPM stem from multi-lineage progenitors, such as the *Scl/Tal1*-positive hemangioblasts that contribute to endothelial and hematopoietic lineages (Choi et al., 1998; Murray, 1932; Sabin, 1917; Vogeli et al., 2006). Our single-cell transcriptomics analysis of the early *drl* reporter-expressing LPM now captured a surprising heterogeneity within the forming LPM already at the end of zebrafish gastrulation, including putative cardiopharyngeal, endothelial/hematopoietic, and here defined mesothelial progenitor clusters (**Fig. 2**). These data complement and expand prior scRNA-seq in mouse, zebrafish, and tunicate embryos that captured numerous mesendodermal and mesodermal cell fate territories at various developmental stages (Farrell et al., 2018; Mohammed et al., 2017; Pijuan-Sala et al., 2019; Scialdone et al., 2016; Wagner et al., 2018; Wang et al., 2019a). With *wt1a/b* expression appearing later than *hand2* and seemingly restricted to the visceral mesothelial layers (**Fig. 3**), *hand2* provides the earliest and most complete marker for mesothelial progenitors in zebrafish and possibly beyond. Our findings are applicable to *in vitro* differentiation protocols for multilineage organoids with desirable stromal components, such as to guide pluripotent stem cell differentiation into representative cell types of the gastro-intestinal tract. Further, our work provides a framework to interpret cell types deemed mesenchymal in numerous systems.

Our results provide several new insights into mesothelial progenitor biology. Combined expression of *hand2* with *sfrp5, meis3*, and *foxh1* potentially defines a rudimentary mesothelial progenitor signature in the early LPM (**Fig. 2E-K; Fig. S4**). Individually, these genes have all previously been linked to mesothelial biology. *sfrp5* expression has been reported in mesenchymal cells around internal organs in zebrafish and in the foregut-surrounding epithelium in *Xenopus* (Li et al., 2008; Stuckenholz et al., 2013). *meis3* expression has been linked to mesenchymal cells lining the developing intestine and to enteric neuron migration in zebrafish (Reichenbach et al., 2008; Uribe and Bronner, 2015); we note that the potentially redundant *meis2a/b* are also expressed in the early LPM and mesothelial progenitors (**Fig. S4**). During segmentation stages, *foxh1* is involved in controlling left-right asymmetry in response to Nodal signaling in a lateral LPM domain of so-far unassigned fate (Slagle et al., 2011; Tavares et al., 2007); our data now proposes that this activity of *foxh1*, and possibly the response to left-right cues, is confined to the *hand2*-positive mesothelial progenitors (**Fig. 2H**). We also observed that the PGCs migrate already from gastrulation stages associated with the mesothelium-primed LPM that establishes their ventral migration border (**Fig. 4**). Recent scRNA-seq studies of early mouse embryos have uncovered *Hand2-*expressing cell clusters that were deemed mesenchymal and that we now likely recognize as mesothelial progenitors by their complement of LPM-associated transcripts (**Fig. S5**) (Han et al., 2020; Pijuan-Sala et al., 2020). Altogether, these observations connect the coelomic epithelium progenitors to key aspects of body plan organization already in early development. The exact composition of a mesothelial progenitor-controlling program and its evolutionary conservation beyond the genes uncovered here warrants further attention.

Expanding previous loss-of-function analyses that documented *Hand* gene contribution to heart and forelimb development (Firulli et al., 1998; Firulli et al., 2010; Osterwalder et al., 2014; Yelon et al., 2000), we here add that mesothelial progenitors fail to properly migrate in *hand2*-mutant zebrafish, resulting in ventral herniation of the yolk at 3 dpf due to incomplete closure of the ventral body wall (**Fig. 6**). How loss of *hand2* leads to dysregulated migration of the mesothelial progenitors warrants future efforts. In congenital malformations affecting the body wall including omphalocele and gastroschisis, the future ventral abdominal body wall fails to form properly, resulting in internal organ protrusion and herniation of abdominal muscles, among other phenotypes (Boylan et al., 2020; Brewer and Williams, 2004; Sadler and Feldkamp, 2008). Among the candidate factors involved in this process is TGF-β signaling that has been repeatedly associated with Hand2 function and midline migration (Gays et al., 2017; Laurent et al., 2017; Yin et al., 2010), as well as Six4 and Six5 (Takahashi et al., 2018). Further, Hand2 regulates extracellular matrix components including repression of Fibronectin production, disruption of which lead to migration defects in the heart and prospective visceral mesothelium (Garavito-Aguilar et al., 2010; Yin et al., 2010). Altogether, our data indicate that *hand2* is not only active early on during mesothelium emergence but also functionally contributes to correct mesothelium formation in zebrafish.

Asbestos-induced mesothelioma remains a global health challenge (Carbone et al., 2019; Felley-Bosco and Macfarlane, 2018). While asbestos fibers have been causally linked to pleural mesothelioma (Wagner et al., 1962), the transformative mechanism and the exact cell of origin await clarification. Different patient stratifications have been assigned to mesothelioma with biphasic, epithelioid, or sarcomatoid tumors based on histopathology (Carbone et al., 2019). Nonetheless, only a few genetic markers are known, including YAP activity and altered BAP1 expression, and their role in mesothelioma formation and maintenance remains unclear (Carbone et al., 2019; Hmeljak et al., 2018). Work in mouse models has uncovered activation of stemness-related signaling pathways and an increase in *Msln*-expressing cells upon asbestos exposure (Rehrauer et al., 2018). Human epithelioid mesothelioma show upregulated expression of coelomic epithelium-associated genes, including of *WT1* (Bozzi et al., 2016; de Reyniès et al., 2014). Notably, the non-coding RNA *Fendrr* (*Fetal-lethal noncoding*) is strongly upregulated in mouse and human epithelioid mesothelioma and shares a bidirectional promoter with the LPM transcription factor gene *FOXF1* (Bueno et al., 2016; Felley-Bosco and Rehrauer, 2018; Grote et al., 2013). These observations indicate that at least some populations of mesothelioma cells feature a gene expression profile reminiscent of late developmental markers for the coelomic epithelium.

Aberrant reactivation of early developmental programs is increasingly recognized as contributor to tumorigenesis (Aiello and Stanger, 2016; Kaufman et al., 2016; Pomerantz et al., 2020). Extrapolating from our developmental findings in zebrafish, our data propose that mesothelioma might reactivate LPM-specific factors involved in initial mesothelial progenitor formation: in both mouse and human mesothelioma, we found the expression of Hand2 and additional transcription factor genes associated with early mesothelium-primed LPM (**Fig. 2, 7**). Notably, crocidolite-exposed tissue showed intermediate expression levels for mesothelial progenitor genes, further supporting a model wherein mesothelioma formation involves a temporal re-activation of LPM-expressed genes deployed in initial mesothelium development (**Fig. 7**). While more work is warranted to establish whether and which LPM transcription factors contribute to mesothelioma formation and maintenance, our findings provide several mRNA transcript and protein expression assays to potentially further stratify mesothelioma in the clinic. We speculate that the malignant transformation of adult mesothelium, and in particular Hand2, endows altered cells with features that recapitulate the migration, proliferation, and stemness that distinguished mesothelial progenitor cells already during their initial development in the LPM.

## Materials and Methods

### Zebrafish husbandry

Zebrafish (*Danio rerio*) husbandry and experiments were performed according to the European Communities Council Directive (86/609/EEC), the recommendations of the Swiss authorities (Animal Protection Ordinance), and according to IACUC regulations at the University of Colorado School of Medicine, Anschutz Medical Campus. Protocols and experiments were performed as approved by the cantonal veterinary office of the Canton Zurich (Kantonales Veterinäramt, permit no. 150, TV4209), in accordance with the European Union (EU) directive 2011/63/EU as well as the German Animal Welfare Act, and as by the IACUC at the University of Colorado School of Medicine, Anschutz Medical Campus (protocol no. 00370). All zebrafish were raised, kept, and handled according to established protocols (Westerfield, 2007) if not noted otherwise.

### Transgenic zebrafish lines and transgene activity

Established transgenic and mutant lines used in this study includes *Tg(drl:EGFP)* (Mosimann et al., 2015), *Tg(drl:mCherry*^*zh705*^) (Sánchez-Iranzo et al., 2018b), *Tg(hand2:EGFP)* (Kikuchi et al., 2011), Tg(*pax2.1:EGFP*) (Picker et al., 2002), Tg(*lmo2:dsRed2*) (Zhu et al., 2005), Tg(−*6.8kbwt1a:EGFP*) (Bollig et al., 2009), *Tg(wt1b:EGFP), Tg(kop:mCherry-f’-nos3’UTR)* (Blaser et al., 2005), *Tg(drl:creERT2;alpha-crystallin:YFP*) (Mosimann et al., 2015), *Tg(−6.8kbwt1a:creERT2)* (Sánchez-Iranzo et al., 2018a), *Tg(–3.5ubb:loxP-EGFP-loxP-mCherry)* (Mosimann et al., 2011), *Tg(−1.5hsp70:loxP-STOP-loxP-EGFP;alpha-crystallin:Venus*) (Felker et al., 2018), and *hand2* mutants (*han*^*S6*^) (Yelon et al., 2000).

The construct to generate the transgenic line *TgBAC(wt1b:rtTA-p2A-creERT2)*^*cn19*^ (referred to in the text as *wt1b:creERT*) was generated by BAC recombineering using EL250 bacteria (Lee et al., 2001). Fragments were amplified by PCR, adding 50 nucleotide homology arms. First, the iTol2Amp-γ-crystallin:RFP cassette (Sánchez-Iranzo et al., 2018b) was amplified using primers 1. *pTarBAC_HA1_iTol2_F 5’-gcgtaagcggggcacatttcattacctctttctccgcacccgacatagatCCCTGCTCGAGCCGGGCCCAAGTG-3’* and *pTarBAC_HA2_iTol2CrystRFP_R 5’-gcggggcatgactattggcgcgccggatcgatccttaattaagtctactaTCGAGGTCGACGGTATCGATTAAA-3’* and recombined into the backbone of the BAC clone *CH73-186G17* to replace the BAC-derived *loxP* site. Then, the *rtTA-p2A-iCreERT2* cassette was amplified and recombined replacing the *ATG* of the *wt1b* coding sequence, with primers *wt1b_HA1_rtTA_F 5’-gacattttgaactcagatattctagtgttttgcaacccagaaaatccgtcACCATGGTCGACGCCACAACCAT-3*’ and *wt1b_HA2_FRT_R 5’-gcgctcaggtctctgacatccgatcccatcgggccgcacggctctgtcagGGAGGCTACCATGGAGAAG-3’*. Finally, the Kanamycin resistance cassette was removed by inducing expression of Flipase recombinase in the EL250 bacteria. The final *BAC* was purified with the HiPure Midiprep kit (Invitrogen) and injected along with synthetic *Tol2* mRNA into wildtype strain zebrafish embryos. Sequence information and primer details are freely available upon request.

6-18 somite staged (ss) dual-fluorescent embryos were imaged using a Leica SP8 upright confocal microscope with HCX-Apochromat W U-V-I 20x/0.5 water correction objective.

### Mouse transgenic strains

The *Hand1*^*EGFPCreΔNeo/+*^ (Barnes et al., 2010), *Hand2*^*Cre*^ (*dHand-Cre*) (Ruest et al., 2003) and *ROSA26 Reporter* (*R26R*) (Soriano, 1999) mouse strains have all been used as previously described. The *Hand2*^*Cre*^ transgenic strain was produced using a 7.4 kb genomic fragment from the mouse *Hand2* locus immediately upstream of the transcriptional start site. The *Hand1*^*EGFPCreΔNeo/+*^ strain was produced by knocking a *Cre* cassette into the mouse *Hand1* locus.

### Human tumor specimens

Mesothelioma tumor specimens from the previously described collection were used (Oehl et al., 2018). Non-tumor pleural tissue was received from four patients undergoing mesothelioma-unrelated thoracic surgery. The study was approved by the Institutional Ethical Review Board of the University Hospital Zurich under reference number StV 29-2009.

### Zebrafish CreERT2-based lineage tracing

Genetic lineage tracing experiments were performed using the Cre/*loxP* recombination system with 4-OHT-inducible creERT2 transgenic lines. *hsp70l:Switch* reporter zebrafish (Felker et al., 2018) were crossed with *creERT2* drivers *drl:creERT2* (Mosimann et al., 2015), *wt1a:creERT2*, and *wt1b:creERT2*. Lineage-labeling was induced at indicated time points using fresh or pre-heated (65°C for 10 min) 4-Hydroxytamoxifen (4-OHT) (Sigma H7904) stock solutions in DMSO at a final concentration of 10 μM in E3 embryo medium (Felker et al., 2016). To activate the EGFP transcription in *hsp70l:Switch* strains, embryos were incubated at 37°C for 1 hour, 2-3 hours before fixation. Subsequently, the embryos were fixed in 4% paraformaldehyde (PFA) at 4°C overnight, washed and stored in PBS, and processed for imaging.

### Zebrafish histology and sectioning

Transverse sections were performed as previously described (Gays et al., 2017). 4% PFA fixed zebrafish embryos were embedded in 6% low-melting agarose (LMA, Sigma-Aldrich) dissolved in 0.1% Tween-20 (Sigma-Aldrich) in PBS and dissected into 130 μm slices with a vibratome (Leica VT 1000S). For immunostaining, the sections were incubated for 30 minutes in permeabilization buffer (1% BSA (AMRESCO), 1% DMSO (Sigma-Aldrich), and 0.3% Triton X-100 (Sigma-Aldrich) in PBS) and washed with PBT (0.1% Triton X-100 and 1% BSA in PBS). Antibodies were dissolved in PBT and incubated overnight at 4°C (primary antibody) or for 4 hours at room temperature (secondary antibody). Primary antibody used was α-SM22 alpha antibody (ab14106). Secondary antibody used: goat-anti-rabbit 594 (Alexa Fluor, Life Technologies, 1:300). Sections were mounted with DAPI-containing Vectashield (Cat#H-1200, Vector Laboratories). Images were obtained with the Zeiss LSM710 confocal microscope equipped with Plan-Apochromat 40x/1.3 oil DIC M27 or 40x/1.2 immersion correction DIC M27 objectives. Images were adjusted for brightness and cropped using ImageJ/Fiji (Schindelin et al., 2012).

Embryos for whole-mount immunostainings were processed as described above for the sections. Primary antibodies used were anti-Pax2 (GeneTex, GTX128127, 1:250) and anti-Myosin heavy chain (DSHB, MF20, 1:20). Secondary antibodies used were Alexa Fluor conjugates (Invitrogen). Embryos were either imaged on a Leica SP8 upright confocal microscope or Zeiss Z.1 light sheet microscope (see details below).

### Gene expression analysis and *in situ* hybridization (ISH)

First-strand complementary DNA (cDNA) was generated from pooled zebrafish RNA isolated from different developmental stages using SuperScript III first-strand synthesis kit (Invitrogen). DNA templates were generated using first-strand cDNA as a PCR template and the primers as specified for each gene of interest; for *in vitro* transcription initiation, the T7 promoter *5′-TAATACGACTCACTATAGGG-3′* was added to the 5′ ends of reverse primers. Specific primers per gene were *meis3*: forward *5’-TACCACAGCCCACTACCCTCAGC-3’*, reverse *5’-TAATACGACTCACTATAGGGTCAGCAGGATTTGGTGCAGTTG-3’*; *gata5*: forward *5’-CACCATGTATTCGAGCCTGGCTTTATCTTCC-3’*, reverse *5’-TAATACGACTCACTATAGGGTCACGCTTGAGACAGAGCACACC-3’*; *sfrp5*: forward *5’-GAATCACAGCAGAGGATG-3’*, reverse *5’-TAATACGACTCACTATAGGGCATCTGTACTAATG GTCG-3’*; *cdx4*: *5’-CACCATGTATCACCAAGGAGCG-3’*, reverse *5’-TAATACGACTCACTATAGGGTAATCCACAACCCACGCC-3’*. PCR reactions were performed under standard conditions as per manufacturer’s protocol using Phusion high-fidelity DNA polymerase (Thermo Fisher Scientific). RNA probes were generated via overnight incubation at 37°C using 20 U/µL T7 RNA polymerase (Roche) and digoxigenin (DIG)-labeled dNTPs (Roche) as per manufacturer’s protocol. The resulting RNA was precipitated in lithium chloride and EtOH. Wildtype strain zebrafish embryos were fixed in 4% PFA in PBS overnight at 4°C, dechorionated, transferred into 100% MeOH, and stored at −20°C. ISH of whole-mount zebrafish embryos was performed essentially as per standard protocols (Thisse and Thisse, 2008).

### RNAscope assay

The RNAscope assays were carried out based on a previously published protocol (Gross-Thebing et al., 2014) and the manufacturer’s instruction (Advanced Cell Diagnostics). Wildtype strain or transgenic zebrafish embryos of tailbud stage until early somitogenesis were fixed in 4% PFA in PBS for 1 h at RT or overnight at 4°C, hand dechorionation, and transferred into 100% MeOH and stored at −20°C. Protease digestion of embryos using Protease III was performed for 20 min at RT followed by rinsing the embryos three times in 0.01% PBS-T (0.01% Tween-20). Target probe hybridization was performed overnight at 40°C. Target probes were designed by Advanced Cell Diagnostics for *hand2, foxh1*, and *meis3*. For RNA detection, incubation with the different amplifier solutions was performed in an incubator at 40°C. After each hybridization step, the embryos were washed three times with 0.2x SSCT for 15 min. The embryos were then incubated with ACD’s DAPI solution overnight at 4°C. Prior to imaging, embryos were rinsed in 0.01% PBT.

### Tumor RNA extraction, cDNA synthesis, and qPCR

RNA extraction, cDNA synthesis, and qPCR were performed as previously described (Thurneysen et al., 2009). The expression of *HAND2* was determined using the following primers: forward *5’-CAGGACTCAGAGCATCAACAG-3’* and reverse *5’-TCCATGAGGTAGGCGATGTA-3’*. Normalization was performed based on the expression of *Histone H3* detected with primers *5’-GGTAAAGCACCCAGGAAGCA-3’* and *5’-CCTCCAGTAGAGGGCGCAC-3’* (Andre and Felley-Bosco, 2003). Data were analyzed using Mann-Whitney test.

### Tumor immunohistochemistry

Immunohistochemistry was performed as previously described (Frei et al., 2011) using anti-HAND2 antibody (A-12, Sc-398167, 1:100) and secondary universal antibody (Vectastain, Vector laboratories Inc., Burlingame, CA, USA), according to manufacturer instructions. Control staining was performed by omission of the primary antibody.

### Light sheet sample preparation, microscopy, and image analysis

Embryos used for long-term imaging or imaging after 1 dpf were treated with 0.003% 1-phenyl-2-thiourea (PTU, Sigma-Aldrich) to prevent melanin pigment formation.

#### Panoramic SPIM for *hand2:EGFP;drl:mCherry* imaging during somitogenesis

At 50% epiboly, embryos in the chorion were mounted in an FEP tube into 1% LMA. Up to 6 embryos were positioned on top of each other with a minimum gap between them. The FEP tube was mounted in the microscope imaging chamber filled with E3 medium. Time-lapse acquisition was performed with a four-lens SPIM set up by a standardized image acquisition pipeline (Schmid et al., 2013). The microscope was equipped with four identical Olympus UMPlanFL N 10x/0.3 NA detection objectives. The subsequent real-time image processing, registration of time points, and 2D map (Mercator) projections were performed with previously described and published Fiji scripts (Schmid et al., 2013). A Z-stack was obtained from every embryo with an interval of 2 min for a period of 14–17 h. Images were processed using ImageJ/Fiji.

#### Multidirectional SPIM (mSPIM) for long-term *hand2:EGFP;drl:mCherry* imaging

To prevent movement of zebrafish embryos during imaging, we injected 30 pg of α-bungarotoxin mRNA (Swinburne et al., 2015) at the 1-cell stage. At the 20 ss, embryos that showed no sign of muscular contractions were selected and mounted in FEP tubes filled with 0.1% LMA. A detailed mounting protocol can be found in Daetwyler et al., 2019. Long-term time-lapse imaging was performed using a custom multidirectional SPIM (mSPIM) setup with multi-sample capacity (Daetwyler et al., 2019). The microscope was equipped with two Zeiss 10x/0.2 NA illumination objectives and an Olympus UMPlanFL N 10x/0.3 NA detection objective. The embryos were imaged every 20 min for up to 4 days starting around 20 ss from 4 different angles. Custom-made data processing tools were used as previously described (Daetwyler et al., 2019) to stitch the individual volumes per angle. For successful fusion and deconvolution of the 4 individual angles, we manually registered them, and then applied the Fiji “Multiview Reconstruction” and “BigStitcher” plugins (Hörl et al., 2019; Preibisch et al., 2010; Preibisch et al., 2014) for precise registration and deconvolution. ImageJ/Fiji and Imaris (Bitplane AG, Zurich, Switzerland) were used for visualization.

#### Zeiss Z.1

The Zeiss Z.1 microscope equipped with a Zeiss W Plan-Apochromat 20x/0.5 NA objective was used for all other light sheet microscopy and accordingly mentioned in the figure legends. Embryos were either within or out of the chorion embedded in 1% LMA in respectively a 50-or 20-µL glass capillary. Alive embryos older than 16 ss were additionally mounted with 0.016% ethyl 3-aminobenzoate methanesulfonate salt (Tricaine, Sigma) in the LMA and added to the E3 medium to prevent movement during imaging. For live imaging of the heart, the heartbeat was stopped with 30 mM 2,3-butanedione monoxime (BDM, Sigma). Images were processed using ImageJ/Fiji and Imaris.

#### Map projections *drl:EGFP;kop:mCherry-F-actin-nanos-3’UTR*

All pre-processing steps and map projection were performed using custom-developed algorithms in MATLAB. The overall size and shape of the fused embryo were first determined in an image that was binned four times to increase the processing speed. The binned image was binarized using an adaptive threshold (imbinarize function with a sensitivity of 0.4) and the isosurface of the resulting image was determined. A sphere fit was then performed using linear least squares with the ellipsoid_fit function from Mathworks (https://ch.mathworks.com/matlabcentral/fileexchange/24693-ellipsoid-fit). The center and radius of the sphere were then used for performing Mercator map projection of the fused images.

For each fused image, a multi-layered projection was performed by generating concentric circles with different radii (step size: 2 µm). This allowed for unwrapping different layers of the 3D fused embryo onto 2D surfaces. For each layer of the map projection, a dummy image in polar coordinates was first generated that extended from −90° to +90° in latitude and 0° to 360° in longitude. Using Mercator projection formulas, the latitudes and longitudes that correspond to each position in the projected map were then determined. The cartesian (x,y,z) positions that correspond to each of these (latitude, longitude) points in the projected map was obtained using standard spherical to cartesian coordinate system conversion formulas. Once this was obtained, a direct mapping of the pixel values corresponding to an x,y,z position in the fused image onto the projected map was performed yielding the map-projected image.

### Single-cell RNA sequencing of LPM

#### Sample preparation and sorting

Wildtype stain and *drl:mCherry*-positive zebrafish embryos were grown until tailbud stage and chorions were removed by incubating in 1 mg/mL Pronase (Roche), followed by washing in E3 medium. Around 300 embryos were used to generate the sample. The E3 medium was replaced by 1X PBS and the embryos were stored on ice until further processing. The embryos were dissociated using 2 mg/mL collagenase IV (Worthington) in DMEM (high glucose (4.5g/l) and NaHCO3, without L-glutamine and sodium pyruvate, Sigma-Aldrich) and incubated for 5 min in a water bath at 37°C. The embryonic tissues were triturated into a single-cell suspension by pipetting carefully up and down. When the embryos were not yet dissociated sufficiently, they were incubated for another 5 min. Cells were filtered through a 35 μm cell-strainer (Falcon, round-bottom tubes with cell-strainer cap) and centrifuged at 6000 rpm for 30 sec. Cell pellets were resuspended in 1X HBSS (Gibco) containing 2% FBS and subjected to an additional round of centrifugation and resuspension. After washing, the cells were resuspended in 1X PBS.

*drl:mCherry*-positive cells were sorted using a FACS Aria III cell sorter (BD Bioscience). Cells were gated based on size and forward scattering, to exclude debris and doublets. The gating for the negative population was determined based on wildtype tailbud stage embryos. See also Supplementary Figure 2 for gate setting. The SORTseq single-cell RNA-sequencing protocol was carried out as described previously (Muraro et al., 2016). Live mCherry-positive single cells were sorted in four 384-well plates (BioRad) containing 5 μl of CEL-Seq2 primer solution in mineral oil (24 bp polyT stretch, a 4 bp random molecular barcode (UMI), a cell-specific barcode, the 5′Illumina TruSeq small RNA kit adaptor and a T7 promoter), provided by Single Cell Discoveries. After sorting, the plates were immediately placed on ice and stored at −80°C.

Single Cell Discoveries further processed the four plates. In brief, ERCC Spike-in RNA (0.02 μL of 1:50000 dilution) was added to each well before cell lysis with heat shocking. Reverse transcription and second-strand synthesis reagents were dispensed using the Nanodrop II (GC biotech). After generation of cDNA from the original mRNA, all cells from one plate were pooled and the pooled sample was amplified linearly with *in vitro* transcription. The amplified RNA was then reverse transcribed to cDNA using a random hexamer primers (Hashimshony et al., 2016). To generate sequencing libraries, RPI-series index primers were used for library PCR. Libraries were sequenced on an Illumina Nextseq500 using 75 bp paired-end sequencing.

#### Single-cell analysis

The CEL-Seq2 barcoded sequences were obtained from Single Cell Discoveries. STAR v2.5.3a (Dobin et al., 2013) was used to create an index based on the Ensembl *GRCz10.91* genome and annotation, after manually adding *drl:mCherry*. Gene abundances were estimated separately for each plate using zUMIs v0.0.4 (Parekh et al., 2018), retaining cells with at least 100 reads and setting the Hamming distance threshold to 1 for both UMI and cell barcode identification. Finally, the exonic UMI counts for all four plates were merged into a single matrix.

Quality control and filtering were performed using the *scater* R package (McCarthy et al., 2017). Upon removal of genes that were undetected across all cells, we removed cells whose percentage of mitochondrial genes fell beyond 2 Median Absolute Deviations (MADs) of the median. Secondly, features with a count greater than 1 in at least 10 cells were retained for downstream analysis. Finally, we discarded cells measured on plate 4, as it was of overall low quality (low number of counts, high percentage of mitochondrial genes).

Next, we used *Seurat* (Stuart et al., 2019) for clustering and dimension reduction. Clustering was performed using the 2000 most highly variable genes (HVGs) identified via *Seurat*’s *FindVariableFeatures* function with default parameters; clustering and dimension reductions were computed using the first 20 principal components. For clustering, we considered a range of *resolution* parameters (0.2-2.4); downstream analyses were performed on cluster assignments obtained from *resolution* 1.8 (15 subpopulations).

Cluster annotations were performed manually on the basis of canonical markers in conjunction with marker genes identified programmatically with *Seurat*’s *FindAllMarkers* function, and complimentary exploration with *iSEE* (Rue-Albrecht et al., 2018).

For Supplementary Figure 4, the three endoderm clusters were excluded and the remaining clusters manually merged into 6 major subpopulations. For each subpopulation, genes that were differentially expressed (DE) against at 3 others were identified using *scater*’s *findMarkers* function with *pval.type = “some”* (McCarthy et al., 2017). The top 50 DE genes (in terms of effect size) at FDR < 5% and with an average positive log-fold change (*summary.logFC > 0*) were selected for visualization.

#### Software specifications and code availability

All analyses were run in R v4.0.2, with Bioconductor v3.11. Data preprocessing and analysis code are deposited at <u>DOI:10.5281/zenodo.4267898</u>, and available as a browsable *workflowr* website (Blischak et al., 2019). All package versions used throughout this study are captured in the session info provided therein.

### Mesothelioma RNA-sequencing data

#### Mouse model and sample preparation

As previously described, mice were repeatedly intra-peritoneally over a time course of 21 weeks injected with crocidolite or sham-exposed and sacrificed at 33 weeks, 12 weeks after the last crocidolite exposure (Rehrauer et al., 2018) (**Fig. 7**). Tissues, among which mesothelium and tumor masses, were collected from euthanized mice and consecutively processed for histopathological diagnosis and analysis and for RNA-seq analysis. The mice sacrificed corresponded to three groups: sham, crocidolite-exposed with pre-neoplastic lesions, and mice exposed to crocidolite-bearing tumors.

#### Mouse RNA-seq analysis

The RNA isolation, library generation, and RNA-seq analysis pipelines are previously described (Rehrauer et al., 2018), In this study, we extracted the expression values of genes associated with early LPM and mesothelium development and diagnostic markers (*Msln, Wt1, Bap1, Nf2*). Counts were normalized with *DESeq2*’s *estimateSizeFactors* function (Love et al., 2014); heatmap values correspond to log10-transformed, scaled and centered (*scale* with default parameters) normalized counts. Right-hand side row annotations display (unscaled) log10-transformed count ranges.

#### Human RNA-seq analysis

Analysis of human mesothelioma was performed using TCGA data downloaded from cBioportal (www.cBioportal.org) (Gao et al., 2013). To perform unsupervised clustering analysis on the mRNA of TCGA mesothelioma samples, we used the *complexHeatmap* R package (Gu et al., 2016). Heatmap values correspond to log10-transformed, batch-normalized mRNA counts.

## Acknowledgments

We thank Single Cell Discoveries for executing the single-cell RNA-sequencing, as well as the Center for Microscopy and Image Analysis (ZMB) and the Cytometry Facility (FCF) of the University of Zurich for technical support and equipment. We also thank Dr. Deborah Yelon, Dr. Richard M. White, Dr. Charles K. Kaufman, and the members of the Mosimann lab for critical input on the manuscript. Species schematics in Figure 1 and Figure 6 were adjusted from templates provided by PhyloPic (phylopic.org).

## Author contributions

K.D.P. and C.M. conceived the project and designed the study. K.D.P. performed the zebrafish experiments together with E.C.B. and Z.L. K.D.P., S.N., and A.K. performed and interpreted gene expression analysis in zebrafish embryos. K.D.P. and S.N. prepared and interpreted the scRNA-seq. H.L.C., C.S., and M.D.R. processed and analyzed the scRNA-seq data. S.D. and K.D.P. acquired data with multidirectional SPIM (mSPIM) and processed the data. S.R.N. generated map projection scripts. A.E. and H.S. generated *wt1b:creERT2*. D.E.C and A.B.F. performed mouse lineage tracing experiments. M.R. performed the IHC of mouse and human tumor samples, M.R. and J.K. performed qPCR on mesothelioma human patient samples, E.F.B., M.R., and J.K. analyzed the mouse and human samples, J.K. and R.O. analyzed TCGA data. E.R., N.M., J.H., E.F.B., A.B., M.D.R., and C.M. supervised. K.D.P. compiled the data and K.D.P. and C.M. wrote the manuscript with input from all co-authors, E.F.B wrote sections of the manuscript related to mesothelioma.

## Data and material availability

All sequencing datasets generated for this publication will be deposited on the ArrayExpress database upon publication; all reagents used are available upon request.

Raw scRNA-seq data, intermediate, and metadata files to reproduce all analyses and figures are available at <u>DOI:10.6084/m9.figshare.13221053.v1</u>.

MATLAB image processing codes are available at https://github.com/sundar07/Mesothelium.

## Movies

**Movie 1:** Primordial germ cells are migrating within the drl-positive LPM during gastrulation and early somitogenesis

Multiview time-lapse SPIM (Zeiss Z.1) of a representative embryo expressing *drl:EGFP* and primordial germ cell (PGC) marker *nos-3’UTR:mCherry* from 6 hpf until 14 hpf. Already during gastrulation, the PGCs migrate within the *drl*-expressing mesendoderm and concentrate within the LPM during early somitogenesis. Note the cells that end up more posterior in the most lateral territory of the *drl*-positive LPM.

**Movie 2: *hand2* is expressed in the forming mesothelial layers**

Lateral view of dual-color time-lapse imaging of a *hand2:EGFP;drl:mCherry* embryo from 18 hpf until 82 hpf. *drl:mCherry* drives expression in mainly the circulating blood cells from 24 hpf onwards. EGFP expression is restricted to the mesothelial layers, pectoral fin, pharyngeal arches, and heart (endo-, myocardium).

**Movie 3: *hand2*-expressing cells form the pericardium and visceral and parietal peritoneum**

Zebrafish embryo expressing *hand2:EGFP* imaged from four different angles over a time course from 18 hpf until 82 hpf. Full 3D-rendering shows how the EGFP-expressing cells migrate over the yolk and yolk extension to form the parietal peritoneum and the pericardium. A front of cells with a medial-directed migration pattern forms the visceral peritoneum along the anterior-to-posterior axis.

**Supplementary Figure 1:**
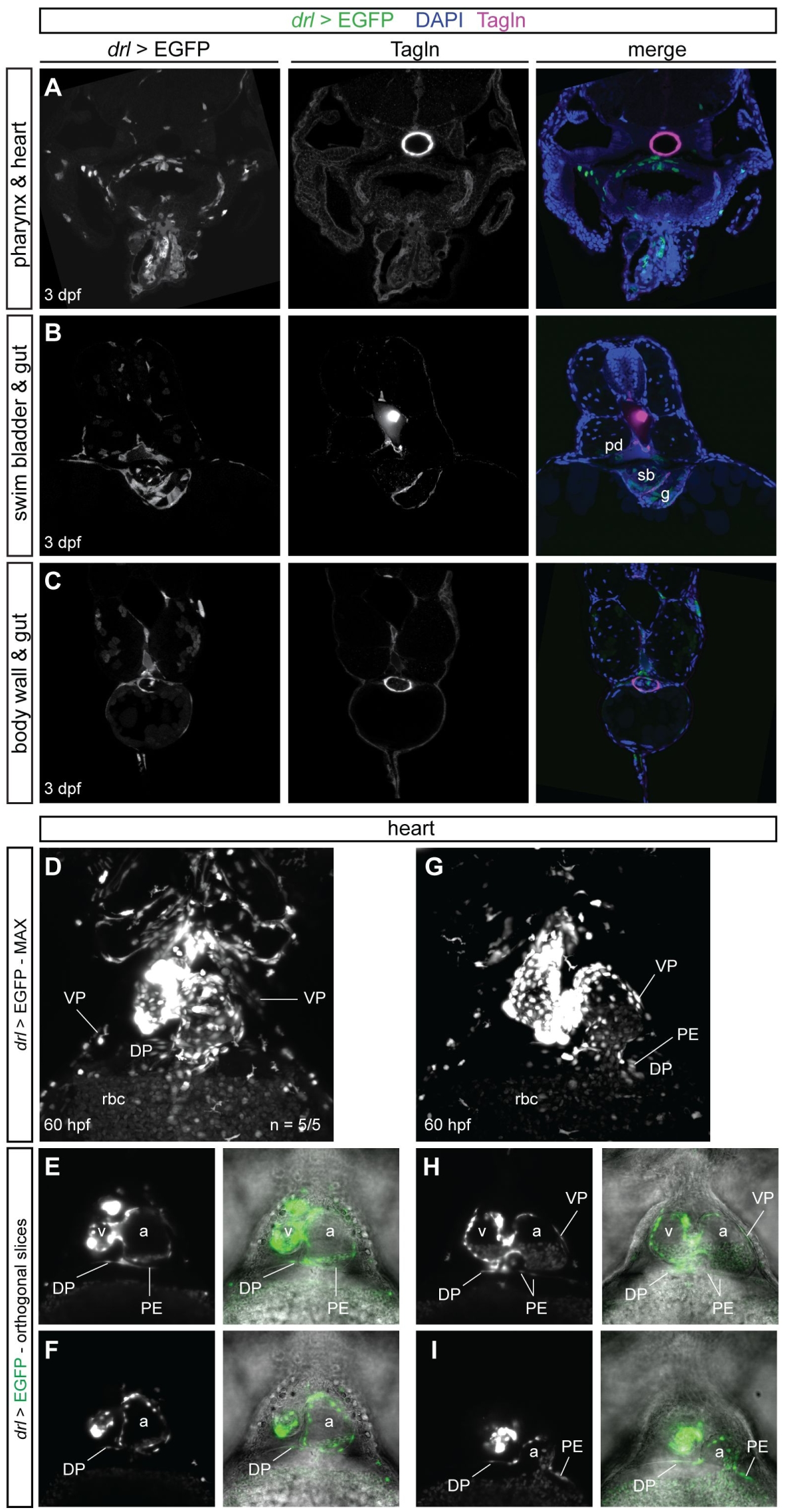
*drl*-derived visceral and parietal mesothelial layers. **(A-C)** Representative transverse sections of LPM lineage-traced *drl:creERT2*; hsp70l:Switch embryos at 3 dpf, co-stained with the smooth muscle marker Transgelin (Tagln). EGFP-expressed was observed in epithelial layers around the pharynx (**A**), swim bladder and gut (**B**), and as part of the body wall (parietal peritoneum) (**C**), in addition to other LPM-derived organs, including vasculature and the kidney tubules. **(D-I)** 4-OHT was administered at tailbud and washed off before 24 hpf. The embryos were heat-shocked at 60 hpf for 1 h and subsequently sorted for EGFP expression. (**D,G**) MAX projections of the heart of two representative *drl*-lineage traced embryos. Prior imaging, the heartbeat was stopped by administering 2,3-butanedione monoxime (BDM) to the embryo water. (**D,F**) EGFP expression in the ventral and dorsal pericardium. (**E,F,H,I**) show orthogonal slices along the Z-axis of the embryos corresponding to (**D**) and (**G**), revealing budding off proepicardial cells. Abbreviations: pronephric ducts (pd), swim bladder (sb), gut (g), ventral pericardium (VP), dorsal pericardium (DP), pro-epicardium (PE), epicardium (EP), red blood cells (rbc), ventricle (v), atrium (a).

**Supplementary Figure 2:**
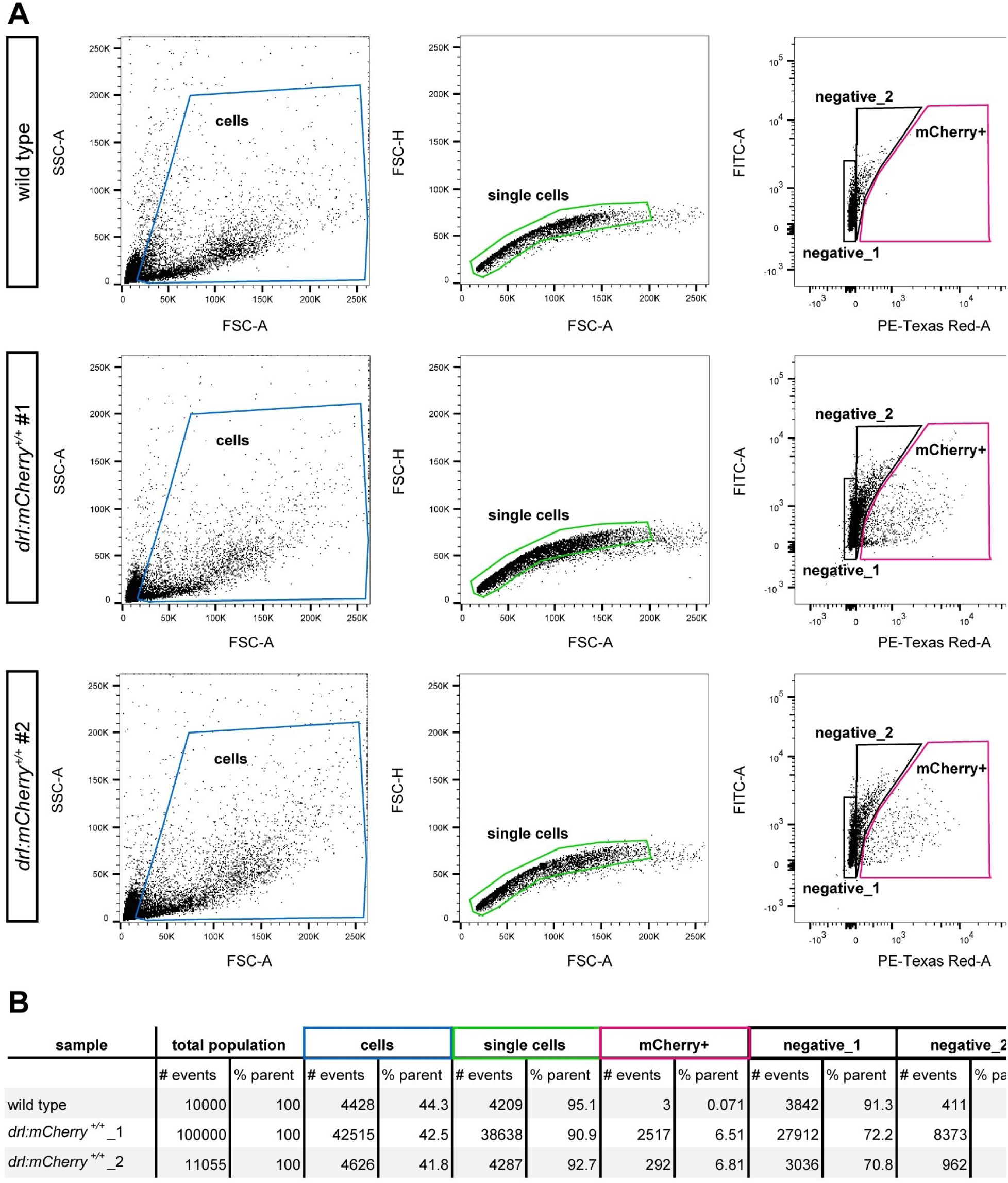
FACS of *drl:mCherry* embryos at tailbud stage. **(A)** FACS plots show gating and sorting strategy for mCherry-positive cells within the single-cell suspension of *drl:mCherry*-dissociated embryos at tailbud stage. **(B)** Table shows the percentage of mCherry-positive cells in the total amount of cells.

**Supplementary Figure 3:**
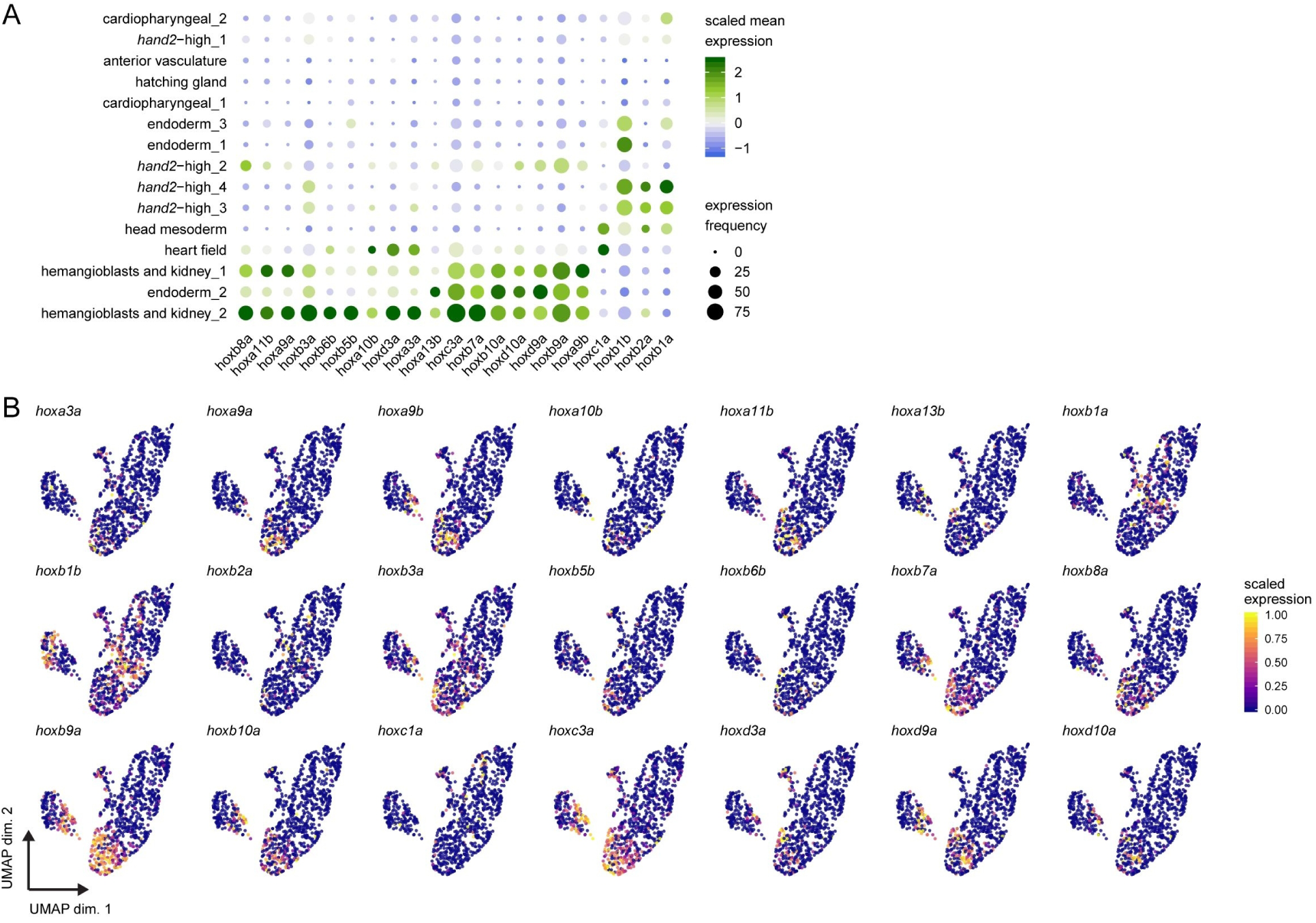
*hox* gene expression across the LPM and endoderm at tailbud stage. **(A)** Dotplot including all *hox* genes expressed throughout the 15 clusters. Dots are colored by column-scaled mean expression (log-transformed library-size-normalized counts), and sized by detection frequency (fraction of cells with non-zero counts); rows and clusters are ordered according to hierarchical clustering of scaled expression values. **(B)** UMAP plots of all the *hox* genes expressed throughout the data set. Cells are colored by scaled expression values using top and bottom 1%-quantiles as boundaries.

**Supplementary Figure 4:**
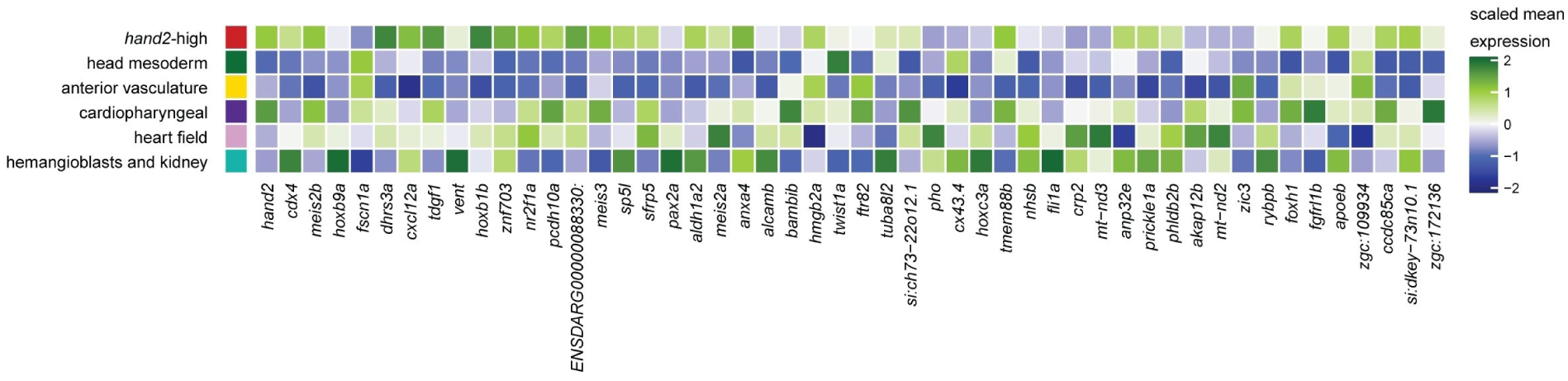
predominant clusters and co-expressed genes within the LPM at tailbud stage. Heatmap of major subpopulation marker genes. Clusters with the same annotation (see **Fig. 2**) were manually merged into 6 ‘super’ clusters: *hand2*-high, head mesoderm, anterior vasculature, cardiopharyngeal, heart field, and hemangioblasts and kidney. Displayed are, for each cluster, the top 50 genes (in terms of effect size) that are differentially expressed against at least half of the remaining clusters (FDR < 5%, average log-fold change > 0); coloring corresponds to scaled mean expression.

**Supplementary Figure 5:**
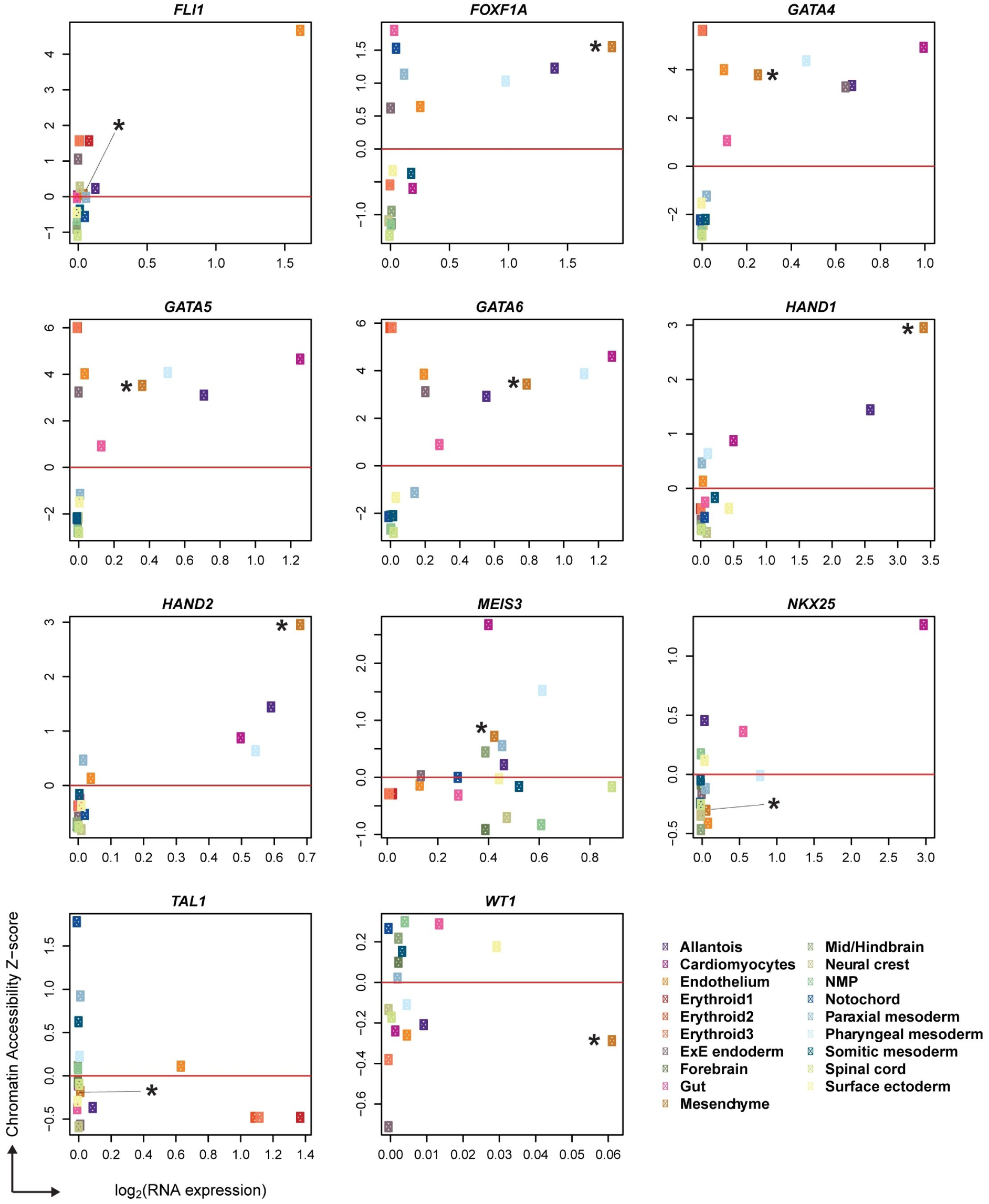
mesothelial progenitors during mouse development as uncovered by co-expression of LPM genes. Single-cell-based gene expression plots as published by Pijuan-Sala et al., 2020. Individual plots depict gene expression distribution of mouse orthologs of LPM-expressed genes, including genes found in zebrafish mesothelial progenitors. Color squares indicate individual organs and cell types, including mesenchyme (asterisks in individual plots). Mouse orthologs of mesothelial progenitor genes as defined in zebrafish including *Hand2, Hand1, GATA4/5/6, Meis3*, and *FoxF1* that is LPM-associated in tetrapods, show high relative expression in mesenchyme clusters, suggesting a mesothelial lineage identity.

**Supplementary Figure 6:**
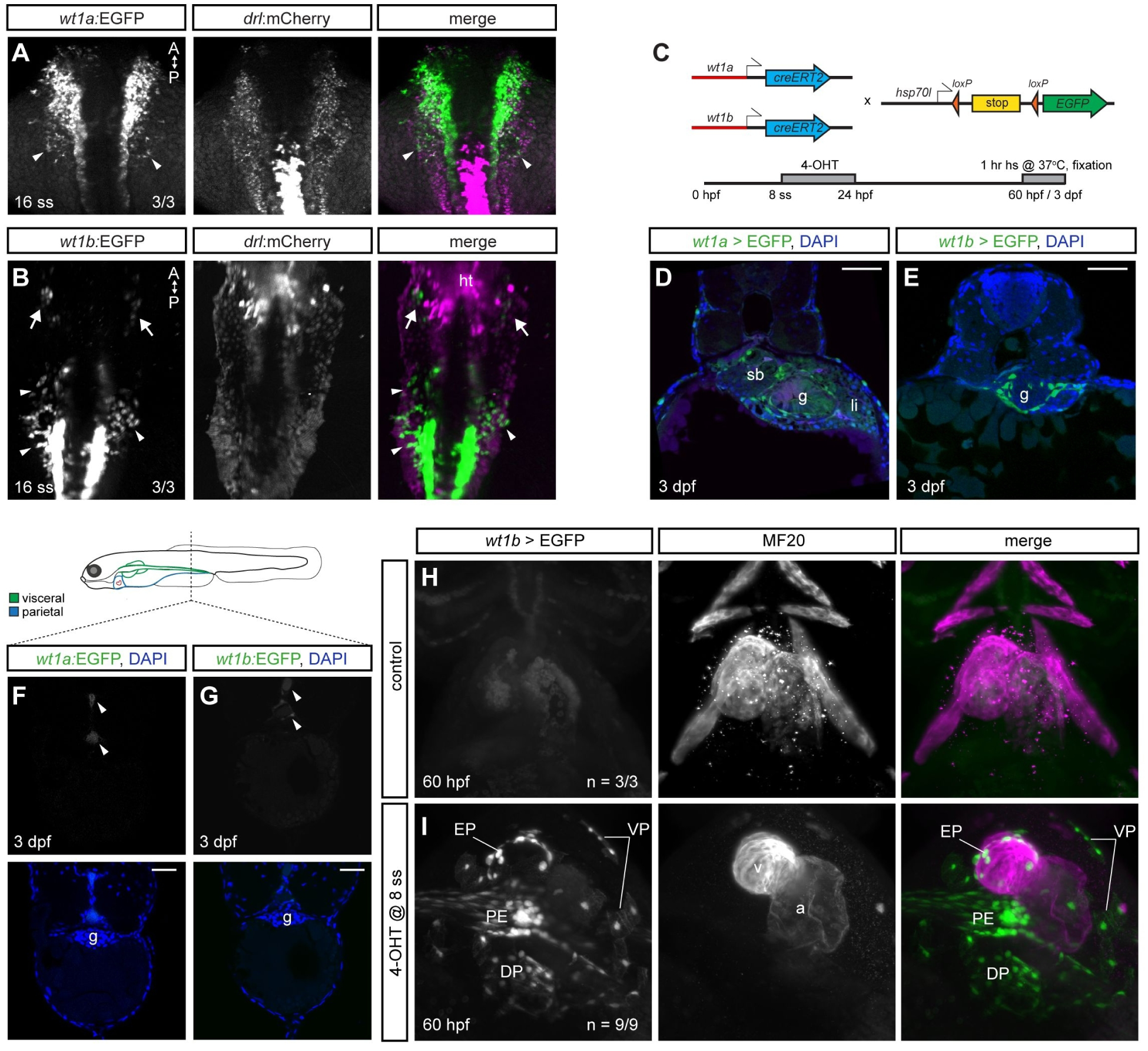
*wt1a* and *wt1b* expression in the zebrafish LPM and in the peritoneal layers. **(A)** Confocal imaging of *wt1a:EGFP;drl:mCherry* embryos at 16 ss, showing EGFP-expression in the developing kidney and a more lateral LPM territory (arrowheads). **(B)** MAX Mercator projection of a *wt1b:EGFP;drl:mCherry* embryo at 16 ss. Note the cells lateral of the developing glomerulus (arrowheads) and in the pericardial progenitor field (arrows). **(C)** Tracing the fate of *wt1a-and wt1b-*derived cells using *wt1a:-* and *wt1b:creERT2* x *hsp70l:Switch*. 4-OHT administered at 8 ss and washed off before 24 hpf. *wt1a/-b>EGFP* indicates lineage labeling. (**D,E**) Transverse sections of *wt1a-* (**D**) and *wt1b-*lineage-traced embryos (**E**), demonstrating labeling of visceral peritoneum. **(F,G)** Posterior trunk transverse sections of *wt1a:-* (**F**) and *wt1b:EGFP*-expressing embryos (**G**), demonstrating absent labeling of the visceral peritoneum around the intestine in more posterior regions and absent labeling of the parietal peritoneum. **(H,I)** SPIM MAX of representative fixed control (n = 3/3) (**H**) and *wt1b-* lineage-traced embryos (n = 9/9) (I) stained with anti-Myosin heavy chain (MF20) in mCherry. Derivatives from *wt1b*-expressing cells are found in ventral and dorsal pericardium, pro-epicardium, and epicardium. Heart tube (ht), swim bladder (sb), gut (g), liver (li), ventral pericardium (VP), dorsal pericardium (DP), pro-epicardium (PE), epicardium (EP), ventricle (v), atrium (a). Nuclei in blue (DAPI). Scale bar (**D-G**) 50 μm.

**Supplementary Figure 7:**
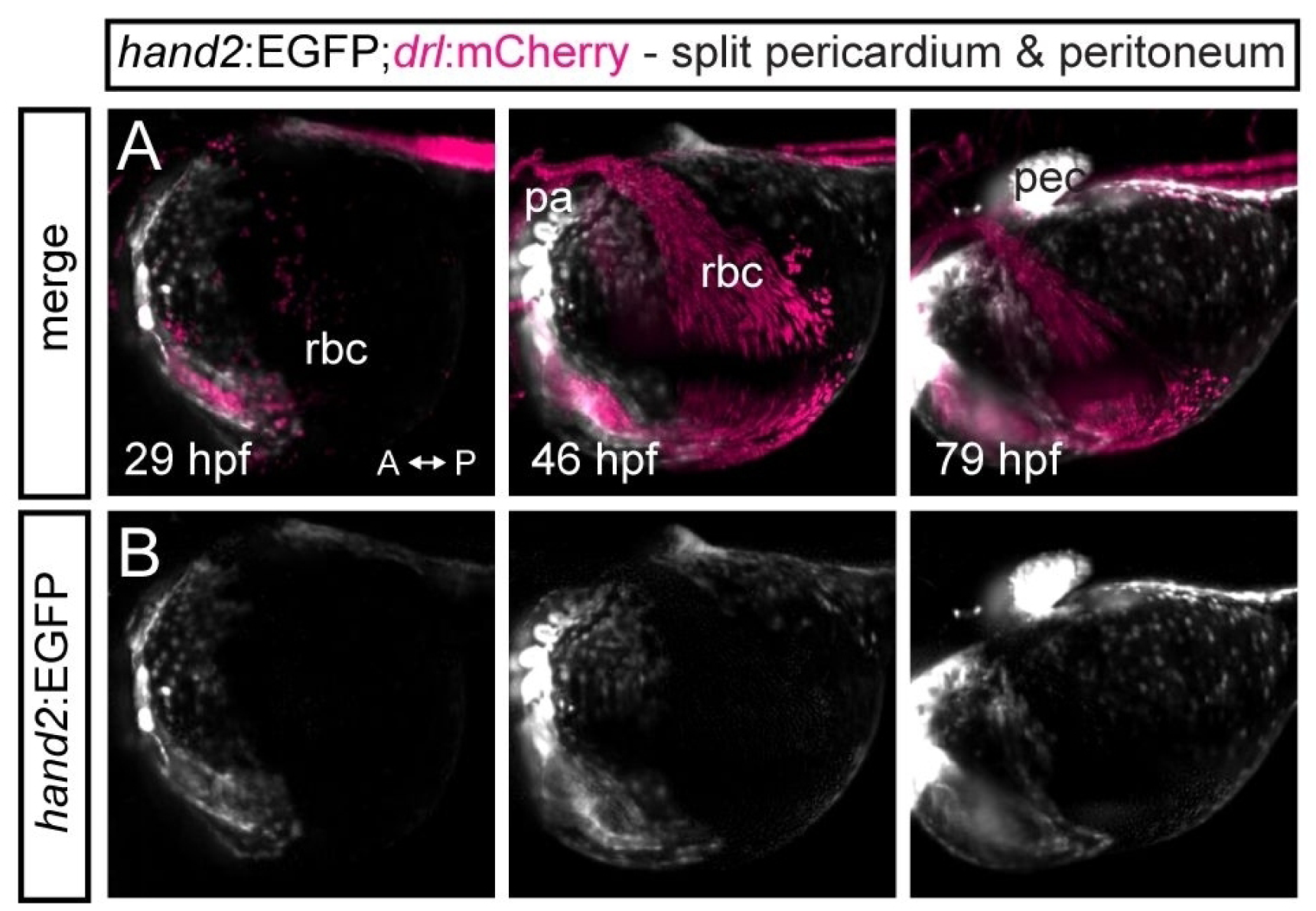
Blood flow over the yolk surface splits pericardium and parietal peritoneum. Maximum intensity projections of the lateral angle of a *hand2:EGFP;drl:mCherry* double-transgenic zebrafish embryo at multiple time points. The imaging shows how the venous blood cells flow over the yolk surface (yolk circulation valley), boarder rostrally by forming pericardium and posterior by the forming parietal peritoneum. The yolk circulation valley is referred to as the duct of Cuvier when a vessel enclosing the blood cells is formed. Abbreviations: red blood cells (rbc), pharyngeal arches (pa), pectoral fin (pec).

## References

Ahn, D. G., Kourakis, M. J., Rohde, L. A., Silver, L. M. and Ho, R. K. (2002). T-box gene tbx5 is essential for formation of the pectoral limb bud. Nature 417, 754–758.

Aiello, N. M. and Stanger, B. Z. (2016). Echoes of the embryo: using the developmental biology toolkit to study cancer. Dis. Model. Mech. 9, 105–14.

Alghamdi, S., Wilm, T. P., Beglinger, S., Boyes, M., Tanton, H., Mutter, F., Allardyce, J., Foisor, V., Middlehurst, B., Carr, L., et al. (2020). Wt1-expressing cells contribute to mesoderm-derived tissues in intestine and mesentery in two distinct phases during murine embryonic development. bioRxiv 2020.08.21.257154.

Andre, M. and Felley-Bosco, E. (2003). Heme oxygenase-1 induction by endogenous nitric oxide: influence of intracellular glutathione. FEBS Lett. 546, 223–227.

Ariza, L., Carmona, R., Cañete, A., Cano, E. and Muñoz-Chápuli, R. (2016). Coelomic epithelium-derived cells in visceral morphogenesis. Dev. Dyn. 245, 307–322.

Asahina, K., Zhou, B., Pu, W. T. and Tsukamoto, H. (2011). Septum transversum-derived mesothelium gives rise to hepatic stellate cells and perivascular mesenchymal cells in developing mouse liver. Hepatology 53, 983– 995.

Barnes, R. M., Firulli, B. A., Conway, S. J., Vincentz, J. W. and Firulli, A. B. (2010). Analysis of the Hand1 cell lineage reveals novel contributions to cardiovascular, neural crest, extra-embryonic, and lateral mesoderm derivatives. Dev. Dyn. 239, 3086–97.

Barnes, R. M., Firulli, B. A., VanDusen, N. J., Morikawa, Y., Conway, S. J., Cserjesi, P., Vincentz, J. W. and Firulli, A. B. (2011). Hand2 loss-of-function in Hand1-expressing cells reveals distinct roles in epicardial and coronary vessel development. Circ. Res. 108, 940–9.

Bianchi, A. B., Mitsunaga, S. I., Cheng, J. Q., Klein, W. M., Jhanwar, S. C., Seizinger, B., Kley, N., Klein-Szanto, A. J. P. and Testa, J. R. (1995). High frequency of inactivating mutations in the neurofibromatosis type 2 gene (NF2) in primary malignant mesotheliomas. Proc. Natl. Acad. Sci. U. S. A.

Blaser, H., Eisenbeiss, S., Neumann, M., Reichman-Fried, M., Thisse, B., Thisse, C. and Raz, E. (2005). Transition from non-motile behaviour to directed migration during early PGC development in zebrafish. J. Cell Sci. 118, 4027–38.

Blischak, J. D., Carbonetto, P. and Stephens, M. (2019). Creating and sharing reproducible research code the workflowr way. F1000Research 8, 1749.

Blondel, V. D., Guillaume, J.-L., Lambiotte, R. and Lefebvre, E. (2008). Fast unfolding of communities in large networks.

Bloomekatz, J., Singh, R., Prall, O. W., Dunn, A. C., Vaughan, M., Loo, C.-S., Harvey, R. P. and Yelon, D. (2017). Platelet-derived growth factor (PDGF) signaling directs cardiomyocyte movement toward the midline during heart tube assembly. Elife 6, e21172.

Bollig, F., Perner, B., Besenbeck, B., Köthe, S., Ebert, C., Taudien, S. and Englert, C. (2009). A highly conserved retinoic acid responsive element controls wt1a expression in the zebrafish pronephros. Development 136, 2883–92.

Bott, M., Brevet, M., Taylor, B. S., Shimizu, S., Ito, T., Wang, L., Creaney, J., Lake, R. A., Zakowski, M. F., Reva, B., et al. (2011). The nuclear deubiquitinase BAP1 is commonly inactivated by somatic mutations and 3p21.1 losses in malignant pleural mesothelioma. In Nature Genetics,.

Boylan, M., Anderson, M. J., Ornitz, D. M. and Lewandoski, M. (2020). The Fgf8 subfamily (Fgf8, Fgf17 and Fgf18) is required for closure of the embryonic ventral body wall. Development.

Bozzi, F., Brich, S., Dagrada, G. P., Negri, T., Conca, E., Cortelazzi, B., Belfiore, A., Perrone, F., Gualeni, A. V., Gloghini, A., et al. (2016). Epithelioid peritoneal mesothelioma: a hybrid phenotype within a mesenchymal-epithelial/epithelial-mesenchymal transition framework. Oncotarget 7, 75503–75517.

Brewer, S. and Williams, T. (2004). Finally, a sense of closure? Animal models of human ventral body wall defects. BioEssays.

Bueno, R., Stawiski, E. W., Goldstein, L. D., Durinck, S., De Rienzo, A., Modrusan, Z., Gnad, F., Nguyen, T. T., Jaiswal, B. S., Chirieac, L. R., et al. (2016). Comprehensive genomic analysis of malignant pleural mesothelioma identifies recurrent mutations, gene fusions and splicing alterations. Nat. Genet.

Cano, E., Carmona, R. and Muñoz-Chápuli, R. (2013). Wt1-expressing progenitors contribute to multiple tissues in the developing lung. Am. J. Physiol. Cell. Mol. Physiol. 305, L322–L332.

Carbone, M., Adusumilli, P. S., Alexander, H. R., Baas, P., Bardelli, F., Bononi, A., Bueno, R., Felley-Bosco, E., Galateau-Salle, F., Jablons, D., et al. (2019). Mesothelioma: Scientific clues for prevention, diagnosis, and therapy. CA. Cancer J. Clin. 69, 402–429.

Carmona, R., Cañete, A., Cano, E., Ariza, L., Rojas, A. and Muñoz-Chápuli, R. (2016). Conditional deletion of WT1 in the septum transversum mesenchyme causes congenital diaphragmatic hernia in mice. Elife 5,.

Carmona, R., Ariza, L., Cano, E., Jiménez-Navarro, M. and Muñoz-Chápuli, R. (2018). Mesothelial-mesenchymal transitions in embryogenesis. Semin. Cell Dev. Biol.

Carney, T. J. and Mosimann, C. (2018). Switch and Trace: Recombinase Genetics in Zebrafish. Trends Genet. 34, 362–378.

Chal, J. and Pourquié, O. (2017). Making muscle: skeletal myogenesis in vivo and in vitro. Development 144,.

Charité, J., Mcfadden, D. G. and Olson, E. N. (2000). The bHLH transcription factor dHAND controls Sonic hedgehog expression and establishment of the zone of polarizing activity during limb development. 2470, 2461–2470.

Chau, Y.-Y., Bandiera, R., Serrels, A., Martínez-Estrada, O. M., Qing, W., Lee, M., Slight, J., Thornburn, A., Berry, R., McHaffie, S., et al. (2014). Visceral and subcutaneous fat have different origins and evidence supports a mesothelial source. Nat. Cell Biol. 16, 367–375.

Chen, Y.-T., Chang, Y.-T., Pan, S.-Y., Chou, Y.-H., Chang, F.-C., Yeh, P.-Y., Liu, Y.-H., Chiang, W.-C., Chen, Y.-M., Wu, K.-D., et al. (2014). Lineage Tracing Reveals Distinctive Fates for Mesothelial Cells and Submesothelial Fibroblasts during Peritoneal Injury. J. Am. Soc. Nephrol. 25, 2847–2858.

Cheng, J. Q., Jhanwar, S. C., Klein, W. M., Bell, D. W., Lee, W.-C., Altomare, D. A., Nobori, T., Olopade, O. I., Buckler, A. J. and Testa, J. R. (1994). p16 Alterations and Deletion Mapping of 9p21–p22 in Malignant Mesothelioma. Cancer Res. 54, 5547–5551.

Chiabrando, J. G., Bonaventura, A., Vecchié, A., Wohlford, G. F., Mauro, A. G., Jordan, J. H., Grizzard, J. D., Montecucco, F., Berrocal, D. H., Brucato, A., et al. (2020). Management of Acute and Recurrent Pericarditis: JACC State-of-the-Art Review. J. Am. Coll. Cardiol.

Choi, K., Kennedy, M., Kazarov, A., Papadimitriou, J. C. and Keller, G. (1998). A common precursor for hematopoietic and endothelial cells. Development 125, 725–732.

Daetwyler, S., Günther, U., Modes, C. D., Harrington, K. and Huisken, J. (2019). Multi-sample SPIM image acquisition, processing and analysis of vascular growth in zebrafish. Development 146, dev173757.

Davidson, A. J. and Zon, L. I. (2004). The “definitive” (and ‘primitive’) guide to zebrafish hematopoiesis. Oncogene 23, 7233–7246.

de Reyniès, A., Jaurand, M.-C., Renier, A., Couchy, G., Hysi, I., Elarouci, N., Galateau-Sallé, F., Copin, M.-C., Hofman, P., Cazes, A., et al. (2014). Molecular Classification of Malignant Pleural Mesothelioma: Identification of a Poor Prognosis Subgroup Linked to the Epithelial-to-Mesenchymal Transition. Clin. Cancer Res. 20, 1323–1334.

Delgado, I., Carrasco, M., Cano, E., Carmona, R., García-Carbonero, R., Marín-Gómez, L. M., Soria, B., Martín, F., Cano, D. A., Muñoz-Chápuli, R., et al. (2014). GATA4 loss in the septum transversum mesenchyme promotes liver fibrosis in mice. Hepatology 59, 2358– 2370.

Dobin, A., Davis, C. A., Schlesinger, F., Drenkow, J., Zaleski, C., Jha, S., Batut, P., Chaisson, M. and Gingeras, T. R. (2013). STAR: ultrafast universal RNA-seq aligner. Bioinformatics 29, 15–21.

Doitsidou, M., Reichman-Fried, M., Stebler, J., Köprunner, M., Dörries, J., Meyer, D., Esguerra, C. V, Leung, T. and Raz, E. (2002). Guidance of primordial germ cell migration by the chemokine SDF-1. Cell 111, 647–59.

Endlich, N., Simon, O., Göpferich, A., Wegner, H., Moeller, M. J., Rumpel, E., Kotb, A. M. and Endlich, K. (2014). Two-Photon Microscopy Reveals Stationary Podocytes in Living Zebra fish Larvae. J. Am. Soc. Nephrol. 25, 681–686.

Farrell, J. A., Wang, Y., Riesenfeld, S. J., Shekhar, K., Regev, A. and Schier, A. F. (2018). Single-cell reconstruction of developmental trajectories during zebrafish embryogenesis. Science 360,.

Felker, A., Nieuwenhuize, S., Dolbois, A., Blazkova, K., Hess, C., Low, L. W. L., Burger, S., Samson, N., Carney, T. J., Bartunek, P., et al. (2016). In Vivo Performance and Properties of Tamoxifen Metabolites for CreERT2 Control. PLoS One 11, e0152989.

Felker, A., Prummel, K. D., Merks, A. M., Mickoleit, M., Brombacher, E. C., Huisken, J., Panáková, D. and Mosimann, C. (2018). Continuous addition of progenitors forms the cardiac ventricle in zebrafish. Nat. Commun. 9,.

Felley-Bosco, E. and Macfarlane, M. (2018). Asbestos: Modern Insights for Toxicology in the Era of Engineered Nanomaterials. American Chemical Society.

Felley-Bosco, E. and Rehrauer, H. (2018). Non-coding transcript heterogeneity in mesothelioma: Insights from asbestos-exposed mice. Int. J. Mol. Sci.

Fernandez-Teran, M., Piedra, M. E., Kathiriya, I. S., Srivastava, D., Rodriguez-Rey, J. C. and Ros, M. A. (2000). Role of dHAND in the anterior-posterior polarization of the limb bud: Implications for the Sonic hedgehog pathway. Development.

Firulli, A. B., McFadden, D. G., Lin, Q., Srivastava, D. and Olson, E. N. (1998). Heart and extra-embryonic mesodermal defects in mouse embryos lacking the bHLH transcription factor Hand1. Nat. Genet. 18, 266–270.

Firulli, B. A., McConville, D. P., Byers III, J. S., Vincentz, J. W., Barnes, R. M. and Firulli, A. B. (2010). Analysis of a Hand1 hypomorphic allele reveals a critical threshold for embryonic viability. Dev. Dyn. 239, 2748–2760.

Frei, C., Opitz, I., Soltermann, A., Fischer, B., Moura, U., Rehrauer, H., Weder, W., Stahel, R. and Felley-Bosco, E. (2011). Pleural mesothelioma side populations have a precursor phenotype. Carcinogenesis.

Funato, N., Chapman, S. L., McKee, M. D., Funato, H., Morris, J. A., Shelton, J. M., Richardson, J. A. and Yanagisawa, H. (2009). Hand2 controls osteoblast differentiation in the branchial arch by inhibiting DNA binding of Runx2. Development 136, 615–25.

Funayama, N., Sato, Y., Matsumoto, K., Ogura, T. and Takahashi, Y. (1999). Coelom formation?: binary decision of the lateral plate mesoderm is controlled by the ectoderm. 4138, 4129–4138.

Galli, A., Robay, D., Osterwalder, M., Bao, X., Bénazet, J. D., Tariq, M., Paro, R., Mackem, S. and Zeller, R. (2010). Distinct roles of Hand2 in initiating polarity and posterior Shh expression during the onset of mouse limb bud development. PLoS Genet.

Gao, J., Aksoy, B. A., Dogrusoz, U., Dresdner, G., Gross, B., Sumer, S. O., Sun, Y., Jacobsen, A., Sinha, R., Larsson, E., et al. (2013). Integrative analysis of complex cancer genomics and clinical profiles using the cBioPortal. Sci. Signal. 6, pl1.

Garavito-Aguilar, Z. V., Riley, H. E. and Yelon, D. (2010). Hand2 ensures an appropriate environment for cardiac fusion by limiting Fibronectin function. Development 137,.

Gays, D., Hess, C., Camporeale, A., Ala, U., Provero, P., Mosimann, C. and Santoro, M. M. (2017). An exclusive cellular and molecular network governs intestinal smooth muscle cell differentiation in vertebrates. Development 144, 464–478.

Gibb, N., Lazic, S., Yuan, X., Deshwar, A. R., Leslie, M., Wilson, M. D. and Scott, I. C. (2018). Hey2 regulates the size of the cardiac progenitor pool during vertebrate heart development. Development dev. 167510.

Grimaldi, C. and Raz, E. (2020). Germ cell migration—Evolutionary issues and current understanding. Semin. Cell Dev. Biol. 100, 152–159.

Gross-Thebing, T., Paksa, A. and Raz, E. (2014). Simultaneous high-resolution detection of multiple transcripts combined with localization of proteins in whole-mount embryos. BMC Biol. 12, 55.

Grote, P., Wittler, L., Hendrix, D., Koch, F., Währisch, S., Beisaw, A., Macura, K., Bläss, G., Kellis, M., Werber, M., et al. (2013). The tissue-specific lncRNA Fendrr is an essential regulator of heart and body wall development in the mouse. Dev. Cell 24, 206–14.

Gu, Z., Eils, R. and Schlesner, M. (2016). Complex heatmaps reveal patterns and correlations in multidimensional genomic data. Bioinformatics 32, 2847–2849.

Hamm, M. J., Kirchmaier, B. C. and Herzog, W. (2016). Sema3d controls collective endothelial cell migration by distinct mechanisms via nrp1 and plxnD1.

Han, Z. and Olson, E. N. (2005). Hand is a direct target of Tinman and GATA factors during Drosophila cardiogenesis and hematopoiesis. Development 132, 3525–3536.

Han, L., Chaturvedi, P., Kishimoto, K., Koike, H., Nasr, T., Iwasawa, K., Giesbrecht, K., Witcher, P. C., Eicher, A., Haines, L., et al. (2020). Single cell transcriptomics identifies a signaling network coordinating endoderm and mesoderm diversification during foregut organogenesis. Nat. Commun. 11, 4158.

Hartenstein, V. and Mandal, L. (2006). The blood/vascular system in a phylogenetic perspective. BioEssays 28, 1203–1210.

Hashimshony, T., Senderovich, N., Avital, G., Klochendler, A., de Leeuw, Y., Anavy, L., Gennert, D., Li, S., Livak, K. J., Rozenblatt-Rosen, O., et al. (2016). CEL-Seq2: sensitive highly-multiplexed single-cell RNA-Seq. Genome Biol. 17, 77.

Hinz, T. K. and Heasley, L. E. (2019). Translating mesothelioma molecular genomics and dependencies into precision oncology-based therapies. Semin. Cancer Biol.

Hmeljak, J., Sanchez-Vega, F., Hoadley, K. A., Shih, J., Stewart, C., Heiman, D., Tarpey, P., Danilova, L., Drill, E., Gibb, E. A., et al. (2018). Integrative Molecular Characterization of Malignant Pleural Mesothelioma. Cancer Discov. 8, 1548–1565.

Hogan, B. M., Pase, L., Hall, N. E. and Lieschke, G. J. (2006). Characterisation of duplicate zinc finger like 2 erythroid precursor genes in zebrafish. Dev. Genes Evol. 216, 523–529.

Hörl, D., Rojas Rusak, F., Preusser, F., Tillberg, P., Randel, N., Chhetri, R. K., Cardona, A., Keller, P. J., Harz, H., Leonhardt, H., et al. (2019). BigStitcher: reconstructing high-resolution image datasets of cleared and expanded samples. Nat. Methods 16, 870–874.

Howe, K., Clark, M. D., Torroja, C. F., Torrance, J., Berthelot, C., Muffato, M., Collins, J. E., Humphray, S., McLaren, K., Matthews, L., et al. (2013). The zebrafish reference genome sequence and its relationship to the human genome. Nature 496, 498–503.

Jiang, Y., Drysdale, T. A. and Evans, T. (1999). A Role for GATA-4/5/6 in the Regulation of Nkx2.5 Expression with Implications for Patterning of the Precardiac Field. Dev. Biol. 216, 57–71.

John, L. B., Trengove, M. C., Fraser, F. W., Yoong, S. H. and Ward, A. C. (2013). Pegasus, the “atypical” Ikaros family member, influences left-right asymmetry and regulates pitx2 expression. Dev. Biol.

Kaufman, C. K., Mosimann, C., Fan, Z. P., Yang, S., Thomas, A. J., Ablain, J., Tan, J. L., Fogley, R. D., van Rooijen, E., Hagedorn, E. J., et al. (2016). A zebrafish melanoma model reveals emergence of neural crest identity during melanoma initiation. Science 351, aad2197.

Kikuchi, K., Holdway, J. E., Major, R. J., Blum, N., Dahn, R. D., Begemann, G. and Poss, K. D. (2011). Short Article Retinoic Acid Production by Endocardium and Epicardium Is an Injury Response Essential for Zebrafish Heart Regeneration. Dev. Cell 20, 397–404.

Kishimoto, K., Furukawa, K. T., Luz-Madrigal, A., Yamaoka, A., Matsuoka, C., Habu, M., Alev, C., Zorn, A. M. and Morimoto, M. (2020). Bidirectional Wnt signaling between endoderm and mesoderm confers tracheal identity in mouse and human cells. Nat. Commun. 11, 4159.

Koopmans, T. and Rinkevich, Y. (2018). Mesothelial to mesenchyme transition as a major developmental and pathological player in trunk organs and their cavities. Commun. Biol. 1, 170.

Laurent, F., Girdziusaite, A., Gamart, J., Barozzi, I., Osterwalder, M., Akiyama, J. A., Lincoln, J., Lopez-Rios, J., Visel, A., Zuniga, A., et al. (2017). HAND2 Target Gene Regulatory Networks Control Atrioventricular Canal and Cardiac Valve Development. Cell Rep. 19, 1602–1613.

Lawson, N. D. and Weinstein, B. M. (2002). In vivo imaging of embryonic vascular development using transgenic zebrafish. Dev. Biol. 248, 307– 18.

Lee, K.-H., Xu, Q. and Breitbart, R. E. (1996). A Newtinman-Related Gene,nkx2.7,Anticipates the Expression ofnkx2.5andnkx2.3in Zebrafish Heart and Pharyngeal Endoderm. Dev. Biol. 180, 722–731.

Lee, E. C., Yu, D., Martinez de Velasco, J., Tessarollo, L., Swing, D. A., Court, D. L., Jenkins, N. A. and Copeland, N. G. (2001). A highly efficient Escherichia coli-based chromosome engineering system adapted for recombinogenic targeting and subcloning of BAC DNA. Genomics 73, 56–65.

Lee, K. Y., Luong, Q., Sharma, R., Dreyfuss, J. M., Ussar, S. and Kahn, C. R. (2019). Developmental and functional heterogeneity of white adipocytes within a single fat depot. EMBO J. 38,.

Lewellis, S. W. and Knaut, H. (2012). Attractive guidance: how the chemokine SDF1/CXCL12 guides different cells to different locations. Semin. Cell Dev. Biol. 23, 333–40.

Li, Y., Rankin, S. A., Sinner, D., Kenny, A. P., Krieg, P. A. and Zorn, A. M. (2008). Sfrp5 coordinates foregut specification and morphogenesis by antagonizing both canonical and noncanonical Wnt11 signaling. Genes Dev.

Love, M. I., Huber, W. and Anders, S. (2014). Moderated estimation of fold change and dispersion for RNA-seq data with DESeq2. Genome Biol.

Lu, F., Langenbacher, A. and Chen, J.-N. (2017). Tbx20 drives cardiac progenitor formation and cardiomyocyte proliferation in zebrafish. Dev. Biol. 421, 139–148.

Lua, I., James, D., Wang, J., Wang, K. S. and Asahina, K. (2014). Mesodermal mesenchymal cells give rise to myofibroblasts, but not epithelial cells, in mouse liver injury. Hepatology 60, 311–22.

Mahlapuu, M., Ormestad, M., Enerbäck, S. and Carlsson, P. (2001). The forkhead transcription factor Foxf1 is required for differentiation of extra-embryonic and lateral plate mesoderm. Development 128, 155–66.

Mao, Q., Stinnett, H. K. and Ho, R. K. (2015). Asymmetric cell convergence-driven zebrafish fin bud initiation and pre-pattern requires Tbx5a control of a mesenchymal Fgf signal. Development 142, 4329–39.

Martínez-Estrada, O. M., Lettice, L. A., Essafi, A., Guadix, J. A., Slight, J., Velecela, V., Hall, E., Reichmann, J., Devenney, P. S., Hohenstein, P., et al. (2010). Wt1 is required for cardiovascular progenitor cell formation through transcriptional control of Snail and E-cadherin. Nat. Genet. 42, 89–93.

McCarthy, D. J., Campbell, K. R., Lun, A. T. L. and Wills, Q. F. (2017). Scater: pre-processing, quality control, normalization and visualization of single-cell RNA-seq data in R. Bioinformatics 33, btw777.

Mohammed, H., Hernando-Herraez, I., Savino, A., Scialdone, A., Macaulay, I., Mulas, C., Chandra, T., Voet, T., Dean, W., Nichols, J., et al. (2017). Single-Cell Landscape of Transcriptional Heterogeneity and Cell Fate Decisions during Mouse Early Gastrulation. Cell Rep. 20, 1215–1228.

Monahan-Earley, R., Dvorak, A. M. and Aird, W. C. (2013). Evolutionary origins of the blood vascular system and endothelium. J. Thromb. Haemost. 11 Suppl 1, 46–66.

Mosimann, C., Kaufman, C. K., Li, P., Pugach, E. K., Tamplin, O. J. and Zon, L. I. (2011). Ubiquitous transgene expression and Cre-based recombination driven by the ubiquitin promoter in zebrafish. Development 138, 169–177.

Mosimann, C., Panáková, D., Werdich, A. A., Musso, G., Burger, A., Lawson, K. L., Carr, L. A., Nevis, K. R., Sabeh, M. K., Zhou, Y., et al. (2015). Chamber identity programs drive early functional partitioning of the heart. Nat. Commun. 6,.

Muraro, M. J., Dharmadhikari, G., Grün, D., Groen, N., Dielen, T., Jansen, E., van Gurp, L., Engelse, M. A., Carlotti, F., de Koning, E. J. P., et al. (2016). A Single-Cell Transcriptome Atlas of the Human Pancreas. Cell Syst. 3, 385–394.e3.

Murray, P. D. F. (1932). The Development in vitro of the Blood of the Early Chick Embryo. Proc. R. Soc. B Biol. Sci. 111, 497–521.

Mutsaers, S. E. (2002). Mesothelial cells: their structure, function and role in serosal repair. Respirology 7, 171–91.

Mutsaers, S. E. and Wilkosz, S. (2007). Structure and function of mesothelial cells. Cancer Treat. Res. 134, 1–19.

Mutsaers, S. E., Birnie, K., Lansley, S., Herrick, S. E., Lim, C.-B. and Prêle, C. M. (2015). Mesothelial cells in tissue repair and fibrosis. Front. Pharmacol. 6, 113.

Naganathan, S. R., Popovic, M. and Oates, A. C. (2020). Somite deformations buffer imprecise segment lengths to ensure left-right symmetry. bioRxiv.

Naylor, R. W., Skvarca, L. B., Thisse, C., Thisse, B., Hukriede, N. A. and Davidson, A. J. (2016). BMP and retinoic acid regulate anterior-posterior patterning of the non-axial mesoderm across the dorsal-ventral axis. Nat. Commun. 7, 12197.

Odgerel, C. O., Takahashi, K., Sorahan, T., Driscoll, T., Fitzmaurice, C., Yoko-O, M., Sawanyawisuth, K., Furuya, S., Tanaka, F., Horie, S., et al. (2017). Estimation of the global burden of mesothelioma deaths from incomplete national mortality data. Occup. Environ. Med. 74, 851–858.

Oehl, K., Kresoja-Rakic, J., Opitz, I., Vrugt, B., Weder, W., Stahel, R., Wild, P. and Felley-Bosco, E. (2018). Live-cell mesothelioma biobank to explore mechanisms of tumor progression. Front. Oncol.

Onimaru, K., Shoguchi, E., Kuratani, S. and Tanaka, M. (2011). Development and evolution of the lateral plate mesoderm: comparative analysis of amphioxus and lamprey with implications for the acquisition of paired fins. Dev. Biol. 359, 124–36.

Ormestad, M., Astorga, J. and Carlsson, P. (2004). Differences in the embryonic expression patterns of mouse Foxf1 and -2 match their distinct mutant phenotypes. Dev. Dyn. 229, 328–33.

Osterwalder, M., Speziale, D., Shoukry, M., Mohan, R., Ivanek, R., Kohler, M., Beisel, C., Wen, X., Scales, S. J., Christoffels, V. M., et al. (2014). HAND2 targets define a network of transcriptional regulators that compartmentalize the early limb bud mesenchyme. Dev. Cell 31, 345–57.

Paksa, A., Bandemer, J., Hoeckendorf, B., Razin, N., Tarbashevich, K., Minina, S., Meyen, D., Biundo, A., Leidel, S. A., Peyrieras, N., et al. (2016). Repulsive cues combined with physical barriers and cell-cell adhesion determine progenitor cell positioning during organogenesis. Nat. Commun. 7, 11288.

Parekh, S., Ziegenhain, C., Vieth, B., Enard, W. and Hellmann, I. (2018). zUMIs - A fast and flexible pipeline to process RNA sequencing data with UMIs. Gigascience 7,.

Parenti, R., Perris, R., Vecchio, G. M., Salvatorelli, L., Torrisi, A., Gravina, L. and Magro, G. (2013). Immunohistochemical expression of Wilms’ tumor protein (WT1) in developing human epithelial and mesenchymal tissues. Acta Histochem. 115, 70–75.

Peralta, M., Steed, E., Harlepp, S., González-Rosa, J. M., Monduc, F., Ariza-Cosano, A., Cortés, A., Rayón, T., Gómez-Skarmeta, J.-L., Zapata, A., et al. (2013). Heartbeat-Driven Pericardiac Fluid Forces Contribute to Epicardium Morphogenesis. Curr. Biol. 23, 1726–1735.

Peralta, M., González-Rosa, J. M., Marques, I. J. and Mercader, N. (2014). The Epicardium in the Embryonic and Adult Zebrafish. J. Dev. Biol. 2, 101–116.

Perens, E. A., Garavito-Aguilar, Z. V, Guio-Vega, G. P., Peña, K. T., Schindler, Y. L. and Yelon, D. (2016). Hand2 inhibits kidney specification while promoting vein formation within the posterior mesoderm. Elife 5, e19941.

Perner, B., Bates, T., Naumann, U. and Englert, C. (2016). Function and Regulation of the Wilms’ Tumor Suppressor 1 (WT1) Gene in Fish. In The Wilms ‘Tumor (WT1) Gene (ed. Hastie, N.), pp. 119–128. Humana Press.

Peterkin, T., Gibson, A. and Patient, R. (2009). Development. Development 101, 45–49.

Pfeffer, P. L., Gerster, T., Lun, K., Brand, M. and Busslinger, M. (1998). Characterization of three novel members of the zebrafish Pax2/5/8 family: dependency of Pax5 and Pax8 expression on the Pax2.1 (noi) function. Development 125, 3063–74.

Picker, A., Scholpp, S., Böhli, H., Takeda, H. and Brand, M. (2002). A novel positive transcriptional feedback loop in midbrain-hindbrain boundary development is revealed through analysis of the zebrafish pax2.1 promoter in transgenic lines. Development 129, 3227–39.

Pijuan-Sala, B., Griffiths, J. A., Guibentif, C., Hiscock, T. W., Jawaid, W., Calero-Nieto, F. J., Mulas, C., Ibarra-Soria, X., Tyser, R. C. V., Ho, D. L. L., et al. (2019). A single-cell molecular map of mouse gastrulation and early organogenesis. Nature 1.

Pijuan-Sala, B., Wilson, N. K., Xia, J., Hou, X., Hannah, R. L., Kinston, S., Calero-Nieto, F. J., Poirion, O., Preissl, S., Liu, F., et al. (2020). Single-cell chromatin accessibility maps reveal regulatory programs driving early mouse organogenesis. Nat. Cell Biol. 1–11.

Pogoda, H. M., Solnica-Krezel, L., Driever, W. and Meyer, D. (2000). The zebrafish forkhead transcription factor FoxH1/Fast1 is a modulator of nodal signaling required for organizer formation. Curr. Biol. 10, 1041–9.

Pomerantz, M. M., Qiu, X., Zhu, Y., Takeda, D. Y., Pan, W., Baca, S. C., Gusev, A., Korthauer, K. D., Severson, T. M., Ha, G., et al. (2020). Prostate cancer reactivates developmental epigenomic programs during metastatic progression. Nat. Genet. 52, 790–799.

Preibisch, S., Saalfeld, S., Schindelin, J. and Tomancak, P. (2010). Software for bead-based registration of selective plane illumination microscopy data. Nat. Methods 7, 418–419.

Preibisch, S., Amat, F., Stamataki, E., Sarov, M., Singer, R. H., Myers, E. and Tomancak, P. (2014). Efficient Bayesian-based multiview deconvolution. Nat. Methods 11, 645–8.

Prummel, K. D., Hess, C., Nieuwenhuize, S., Parker, H. J., Rogers, K. W., Kozmikova, I., Racioppi, C., Brombacher, E. C., Czarkwiani, A., Knapp, D., et al. (2019). A conserved regulatory program initiates lateral plate mesoderm emergence across chordates. Nat. Commun. 10, 3857.

Prummel, K. D., Nieuwenhuize, S. and Mosimann, C. (2020). The lateral plate mesoderm. Development 147,.

Quetel, L., Meiller, C., Assie, J.-B., Blum, Y., Imbeaud, S., Montagne, F., Tranchant, R., de Wolf, J., Caruso, S., Copin, M.-C., et al. (2020). Genetic alterations of malignant pleural mesothelioma: association with tumor heterogeneity and overall survival. Mol. Oncol.

Raz, E. (2003). Primordial germ-cell development: the zebrafish perspective. Nat. Rev. Genet. 4, 690–700.

Rehrauer, H., Wu, L., Blum, W., Pecze, L., Henzi, T., Serre-Beinier, V., Aquino, C., Vrugt, B., de Perrot, M., Schwaller, B., et al. (2018). How asbestos drives the tissue towards tumors: YAP activation, macrophage and mesothelial precursor recruitment, RNA editing, and somatic mutations. Oncogene 1.

Reichenbach, B., Delalande, J.-M., Kolmogorova, E., Prier, A., Nguyen, T., Smith, C. M., Holzschuh, J. and Shepherd, I. T. (2008). Endoderm-derived Sonic hedgehog and mesoderm Hand2 expression are required for enteric nervous system development in zebrafish. Dev. Biol. 318, 52–64.

Reiter, J. F., Alexander, J., Rodaway, A., Yelon, D., Patient, R., Holder, N. and Stainier, D. Y. (1999). Gata5 is required for the development of the heart and endoderm in zebrafish. Genes Dev. 13, 2983–95.

Richardson, B. E. and Lehmann, R. (2010). Mechanisms guiding primordial germ cell migration: strategies from different organisms. Nat. Rev. Mol. Cell Biol. 11, 37–49.

Rinkevich, Y., Mori, T., Sahoo, D., Xu, P.-X., Bermingham, J. R. and Weissman, I. L. (2012). Identification and prospective isolation of a mesothelial precursor lineage giving rise to smooth muscle cells and fibroblasts for mammalian internal organs and their vasculature. Nat. Cell Biol. 14, 1251–1260.

Rue-Albrecht, K., Marini, F., Soneson, C. and Lun, A. T. L. (2018). iSEE: Interactive SummarizedExperiment Explorer. F1000Research 7, 741.

Ruest, L.-B., Dager, M., Yanagisawa, H., Charité, J., Hammer, R. E., Olson, E. N., Yanagisawa, M. and Clouthier, D. E. (2003). dHAND-Cre transgenic mice reveal specific potential functions of dHAND during craniofacial development. Dev. Biol. 257, 263–77.

Sabin, F. R. (1917). Preliminary note on the differentiation of angioblasts and the method by which they produce blood-vessels, blood-plasma and red blood-cells as seen in the living chick. 1917. J Hematother Stem Cell Res 11, 5–7.

Sadler, T. W. and Feldkamp, M. L. (2008). The embryology of body wall closure: Relevance to gastroschisis and other ventral body wall defects. Am. J. Med. Genet. Part C Semin. Med. Genet. 148C, 180–185.

Sánchez-Iranzo, H., Galardi-Castilla, M., Sanz-Morejón, A., González-Rosa, J. M., Costa, R., Ernst, A., Sainz de Aja, J., Langa, X. and Mercader, N. (2018a). Transient fibrosis resolves via fibroblast inactivation in the regenerating zebrafish heart. Proc. Natl. Acad. Sci. U. S. A. 115, 4188–4193.

Sánchez-Iranzo, H., Galardi-Castilla, M., Minguillón, C., Sanz-Morejón, A., González-Rosa, J. M. J. M., Felker, A., Ernst, A., Guzmán-Martínez, G., Mosimann, C. and Mercader, N. (2018b). Tbx5a lineage tracing shows cardiomyocyte plasticity during zebrafish heart regeneration. Nat. Commun. 9, 428.

Santoro, M. M., Pesce, G. and Stainier, D. Y. (2009). Characterization of vascular mural cells during zebrafish development. Mech. Dev. 126, 638–649.

Schier, A. F. and Talbot, W. S. (2001). Nodal signaling and the zebrafish organizer. Int. J. Dev. Biol. 45, 289–97.

Schindelin, J., Arganda-Carreras, I., Frise, E., Kaynig, V., Longair, M., Pietzsch, T., Preibisch, S., Rueden, C., Saalfeld, S., Schmid, B., et al. (2012). Fiji: an open-source platform for biological-image analysis. Nat. Methods 9, 676–82.

Schmid, B., Shah, G., Scherf, N., Weber, M., Thierbach, K., Campos, C. P., Roeder, I., Aanstad, P. and Huisken, J. (2013). High-speed panoramic light-sheet microscopy reveals global endodermal cell dynamics. Nat Commun 4, 2207.

Schulte, D. and Geerts, D. (2019). MEIS transcription factors in development and disease. Development 146,.

Scialdone, A., Tanaka, Y., Jawaid, W., Moignard, V., Wilson, N. K., Macaulay, I. C., Marioni, J. C. and Göttgens, B. (2016). Resolving early mesoderm diversification through single-cell expression profiling. Nature 535, 289–293.

Sebo, Z. L., Jeffery, E., Holtrup, B. and Rodeheffer, M. S. (2018). A mesodermal fate map for adipose tissue. Development 145, dev166801.

Sekido, Y., Bader, S., Gazdar, A. F., Minna, J. D., Pass, H. I., Mew, D. J. Y. and Christman, M. F. (1995). Neurofibromatosis Type 2 (NF2) Gene Is Somatically Mutated in Mesothelioma but not in Lung Cancer. Cancer Res.

Sheng, G. (2015). The developmental basis of mesenchymal stem/stromal cells (MSCs). BMC Dev. Biol. 15, 44.

Shih, Y.-H., Zhang, Y., Ding, Y., Ross, C. A., Li, H., Olson, T. M. and Xu, X. (2015). Cardiac Transcriptome and Dilated Cardiomyopathy Genes in Zebrafish. Circ. Cardiovasc. Genet. 8, 261–269.

Shin, M., Nagai, H. and Sheng, G. (2009). Notch mediates Wnt and BMP signals in the early separation of smooth muscle progenitors and blood/endothelial common progenitors. Development 136, 595–603.

Slagle, C. E., Aoki, T. and Burdine, R. D. (2011). Nodal-dependent mesendoderm specification requires the combinatorial activities of FoxH1 and Eomesodermin. PLoS Genet. 7, e1002072.

Soriano, P. (1999). Generalized lacZ expression with the ROSA26 Cre reporter strain. Nat Genet 21, 70–71.

Srivastava, D., Cserjesi, P. and Olson, E. N. (1995). A Subclass of bHLH Proteins Required for Cardiac Morphogenesis. Science (80-.). 270, 1995–1999.

Srivastava, D., Thomas, T., Lin, Q., Kirby, M. L., Brown, D. and Olson, E. N. (1997). Regulation of cardiac mesodermal and neural crest development by the bHLH transcription factor, dHAND. Nat. Genet. 16, 154–60.

Stebler, J., Spieler, D., Slanchev, K., Molyneaux, K. A., Richter, U., Cojocaru, V., Tarabykin, V., Wylie, C., Kessel, M. and Raz, E. (2004). Primordial germ cell migration in the chick and mouse embryo: the role of the chemokine SDF-1/CXCL12. Dev. Biol. 272, 351–361.

Stuart, T., Butler, A., Hoffman, P., Hafemeister, C., Papalexi, E., Mauck, W. M., Hao, Y., Stoeckius, M., Smibert, P. and Satija, R. (2019). Comprehensive Integration of Single-Cell Data. Cell 177, 1888–1902.e21.

Stuckenholz, C., Lu, L., Thakur, P. C., Choi, T.-Y., Shin, D. and Bahary, N. (2013). Sfrp5 modulates both Wnt and BMP signaling and regulates gastrointestinal organogenesis [corrected] in the zebrafish, Danio rerio. PLoS One 8, e62470.

Swinburne, I. A., Mosaliganti, K. R., Green, A. A. and Megason, S. G. (2015). Improved Long-Term Imaging of Embryos with Genetically Encoded α-Bungarotoxin. PLoS One 10, e0134005.

Takahashi, M., Tamura, M., Sato, S. and Kawakami, K. (2018). Mice doubly deficient in Six4 and Six5 show ventral body wall defects reproducing human omphalocele. Dis. Model. Mech. 11,.

Takasato, M. and Little, M. H. (2015). The origin of the mammalian kidney: implications for recreating the kidney in vitro. Development 142, 1937–1947.

Tanaka, M. (2011). Revealing the mechanisms of the rostral shift of pelvic fins among teleost fishes. Evol. Dev. 13, 382–390.

Tanaka, M., Yu, R. and Kurokawa, D. (2015). Anterior migration of lateral plate mesodermal cells during embryogenesis of the pufferfish Takifugu niphobles: insight into the rostral positioning of pelvic fins. J. Anat.

Tate, J. G., Bamford, S., Jubb, H. C., Sondka, Z., Beare, D. M., Bindal, N., Boutselakis, H., Cole, C. G., Creatore, C., Dawson, E., et al. (2019). COSMIC: The Catalogue Of Somatic Mutations In Cancer. Nucleic Acids Res.

Tavares, A. T., Andrade, S., Silva, A. C. and Belo, J. A. (2007). Cerberus is a feedback inhibitor of Nodal asymmetric signaling in the chick embryo. Development 134, 2051–2060.

Technau, U. and Scholz, C. B. (2003). Origin and evolution of endoderm and mesoderm. 47, 531– 539.

Thisse, C. and Thisse, B. (2008). High-resolution in situ hybridization to whole-mount zebrafish embryos. Nat. Protoc. 3, 59–69.

Thurneysen, C., Opitz, I., Kurtz, S., Weder, W., Stahel, R. A. and Felley-Bosco, E. (2009). Functional inactivation of NF2/merlin in human mesothelioma. Lung Cancer 64, 140–147.

Tremblay, M., Sanchez-Ferras, O. and Bouchard, M. (2018). GATA transcription factors in development and disease. Development 145, dev164384.

Tsai, J. M., Sinha, R., Seita, J., Fernhoff, N., Christ, S., Koopmans, T., Krampitz, G. W., McKenna, K. M., Xing, L., Sandholzer, M., et al. (2018). Surgical adhesions in mice are derived from mesothelial cells and can be targeted by antibodies against mesothelial markers. Sci. Transl. Med.

Uribe, R. A. and Bronner, M. E. (2015). Meis3 is required for neural crest invasion of the gut during zebrafish enteric nervous system development. Mol. Biol. Cell 26, 3728–40.

Vogeli, K. M., Jin, S. W., Martin, G. R. and Stainier, D. Y. (2006). A common progenitor for haematopoietic and endothelial lineages in the zebrafish gastrula. Nature 443, 337–339.

Wagner, J. C., Munday, D. E. and Harington, J. S. (1962). Histochemical demonstration of hyaluronic acid in pleural mesotheliomas. J. Pathol. Bacteriol. 84, 73–78.

Wagner, D. E., Weinreb, C., Collins, Z. M., Briggs, J. A., Megason, S. G. and Klein, A. M. (2018). Single-cell mapping of gene expression landscapes and lineage in the zebrafish embryo. Science 360, 981–987.

Walker, C., Rutten, F., Yuan, X., Pass, H., Mew, D. M. and Everitt, J. (1994). Wilms’ tumor suppressor gene expression in rat and human mesothelioma. Cancer Res. 54, 3101–3106.

Wang, W., Niu, X., Stuart, T., Jullian, E., Mauck, W. M., Kelly, R. G., Satija, R. and Christiaen, L. (2019a). A single-cell transcriptional roadmap for cardiopharyngeal fate diversification. Nat. Cell Biol. 21, 674–686.

Wang, H., Holland, P. W. H. and Takahashi, T. (2019b). Gene profiling of head mesoderm in early zebrafish development: insights into the evolution of cranial mesoderm. Evodevo 10, 14.

Warga, R. M. and Nüsslein-Volhard, C. (1999). Origin and development of the zebrafish endoderm. 838, 827–838.

Westerfield, M. (2007). The Zebrafish Book: a guide for the laboratory use of zebrafish (Danio rerio). 5th ed. Eugene: University of Oregon Press.

Winters, N. I., Thomason, R. T. and Bader, D. M. (2012). Identification of a novel developmental mechanism in the generation of mesothelia. Development 139, 2926–2934.

Yap, T. A., Aerts, J. G., Popat, S. and Fennell, D. A. (2017). Novel insights into mesothelioma biology and implications for therapy. Nat. Rev. Cancer.

Yelon, D. and Stainier, D. Y. R. (2005). Hand2 Regulates Epithelial Formation during Myocardial Differentiation. Curr. Biol. 15, 441–446.

Yelon, D., Ticho, B., Halpern, M. E., Ruvinsky, I., Ho, R. K., Silver, L. M. and Stainier, D. Y. R. (2000). The bHLH transcription factor hand2 plays parallel roles in zebrafish heart and pectoral fin development. Development 127, 2573–82.

Yin, C., Kikuchi, K., Hochgreb, T., Poss, K. D. and Stainier, D. Y. R. (2010). Hand2 regulates extracellular matrix remodeling essential for gut-looping morphogenesis in zebrafish. Dev. Cell 18, 973–984.

Zhou, B., Ma, Q., Rajagopal, S., Wu, S. M., Domian, I., Rivera-Feliciano, J., Jiang, D., von Gise, A., Ikeda, S., Chien, K. R., et al. (2008). Epicardial progenitors contribute to the cardiomyocyte lineage in the developing heart. Nature 454, 109–113.

Zhu, H., Traver, D., Davidson, A. J., Dibiase, A., Thisse, C., Thisse, B., Nimer, S. and Zon, L. I. (2005). Regulation of the lmo2 promoter during hematopoietic and vascular development in zebrafish. Dev Biol 281, 256–269.

Zilinski, C. A., Shah, R., Lane, M. E. and Jamrich, M. (2005). Modulation of zebrafish pitx3 expression in the primordia of the pituitary, lens, olfactory epithelium and cranial ganglia by Hedgehog and Nodal signaling. Genesis.

